# Virological characteristics of the SARS-CoV-2 Omicron EG.5.1 variant

**DOI:** 10.1101/2023.10.19.563209

**Authors:** Shuhei Tsujino, Sayaka Deguchi, Tomo Nomai, Miguel Padilla-Blanco, Arnon Plianchaisuk, Lei Wang, MST Monira Begum, Keiya Uriu, Keita Mizuma, Naganori Nao, Isshu Kojima, Tomoya Tsubo, Jingshu Li, Yasufumi Matsumura, Miki Nagao, Yoshitaka Oda, Masumi Tsuda, Yuki Anraku, Shunsuke Kita, Hisano Yajima, Kaori Sasaki-Tabata, Ziyi Guo, Alfredo A Hinay, Kumiko Yoshimatsu, Yuki Yamamoto, Tetsuharu Nagamoto, Hiroyuki Asakura, Mami Nagashima, Kenji Sadamasu, Kazuhisa Yoshimura, Hesham Nasser, Michael Jonathan, Olivia Putri, Yoonjin Kim, Luo Chen, Rigel Suzuki, Tomokazu Tamura, Katsumi Maenaka, The Genotype to Phenotype Japan (G2P-Japan) Consortium, Takashi Irie, Keita Matsuno, Shinya Tanaka, Jumpei Ito, Terumasa Ikeda, Kazuo Takayama, Jiri Zahradnik, Takao Hashiguchi, Takasuke Fukuhara, Kei Sato

## Abstract

In middle-late 2023, a sublineage of SARS-CoV-2 Omicron XBB, EG.5.1 (a progeny of XBB.1.9.2), is spreading rapidly around the world. Here, we performed multiscale investigations to reveal virological features of newly emerging EG.5.1 variant. Our phylogenetic-epidemic dynamics modeling suggested that two hallmark substitutions of EG.5.1, S:F456L and ORF9b:I5T, are critical to the increased viral fitness. Experimental investigations addressing the growth kinetics, sensitivity to clinically available antivirals, fusogenicity and pathogenicity of EG.5.1 suggested that the virological features of EG.5.1 is comparable to that of XBB.1.5. However, the cryo-electron microscopy reveals the structural difference between the spike proteins of EG.5.1 and XBB.1.5. We further assessed the impact of ORF9b:I5T on viral features, but it was almost negligible at least in our experimental setup. Our multiscale investigations provide the knowledge for understanding of the evolution trait of newly emerging pathogenic viruses in the human population.

## Introduction

XBB is a recombinant SARS-CoV-2 Omicron lineage emerged in the summer of 2022^1^. As of October 2023, some XBB sublineages bearing the F486P substitution in the spike protein (S; S:F486P), such as XBB.1.5 and XBB.1.16, have become predominant worldwide (https://nextstrain.org/). Because S:F486P significantly increased pseudovirus infectivity^2^, it is assumed that the spread of F486P-bearing XBB subvariants is attributed to the increased infectivity from S:F486P.

Since July 2023, EG.5.1 (also known as XBB.1.9.2.5.1) has rapidly spread in some Asian and North American countries. On August 9, 2023, the WHO classified EG.5 as a variant of interest^3^. In fact, our recent study showed that EG.5.1 exhibits a greater effective reproduction number (R_e_) compared with XBB.1.5, XBB.1.16, and its parental lineage (XBB.1.9.2)^4^. These observations suggest that EG.5. has the potential to spread globally and outcompete these XBB subvariants.

EG.5.1 bears two evolutionary characteristic mutations, S:F456L and ORF9b:I5T, absent in earlier predominant lineages such as XBB.1.5. These two substitutions convergently occurred in multiple SARS-CoV-2 lineages (https://jbloomlab.github.io/SARS2-mut-fitness/). Importantly, it has been reported that convergent mutations tend to increase viral fitness—the ability of virus to spread in the human population, quantified with the effective reproduction number (R_e_)^5,6^. In fact, we have shown that the S:F456L in EG.5.1 confers resistance to the humoral immunity induced by XBB breakthrough infection (BTI)^4^. This result suggests that S:F456L contributes to the increased viral fitness of EG.5.1 by enhancing immune evasion from the humoral immunity elicited by XBB BTI.

SARS-CoV-2 ORF9b is a viral antagonist that hampers the innate immunity to induce type I interferon (IFN-I) production^7-10^. Of note, the ORF9b:I5T substitution is detected in multiple lineages including XBB.1.9 and EG.5.1 (https://github.com/cov-lineages/pango-designation), which raises a possibility that the ORF9b:I5T has a crucial role in these XBB sublineages. However, the impact of ORF9b:I5T on the characteristics of SARS-CoV-2 variants is not documented yet.

The R_e_ and immune evasive property of SARS-CoV-2 Omicron EG.5.1 variant have been addressed by us^4^ and other groups^11,12^. However, mutations that contribute to the increased viral fitness in EG.5.1 have been unidentified. Moreover, the growth kinetics, sensitivity to clinically available antiviral compounds, fusogenicity and pathogenicity of EG.5.1 remains to be addressed. In this study, we elucidated the virological characteristics of SARS-CoV-2 Omicron EG.5.1 variant.

## Results

### Mutations contributing to the increased viral fitness of EG.5.1

Compared with XBB.1.5, EG.5.1 has two amino acid substitutions in S (S:Q52H and S:F456L) and five substitutions in other proteins (**Figure 1A**). Of these, S:F456L and ORF9b:I5T are documented as convergent substitutions (https://jbloomlab.github.io/SARS2-mut-fitness/). ORF9b:I5T is already present in XBB.1.9, the ancestral lineage of EG.5.1, whereas S:F456L is absent (**Figure 1A**). EG.5.1.1, a major descendant lineage of EG.5.1, has an additional ORF1b:D54N substitution compared with EG.5.1 (**Figure 1A**).

**Figure 1.**
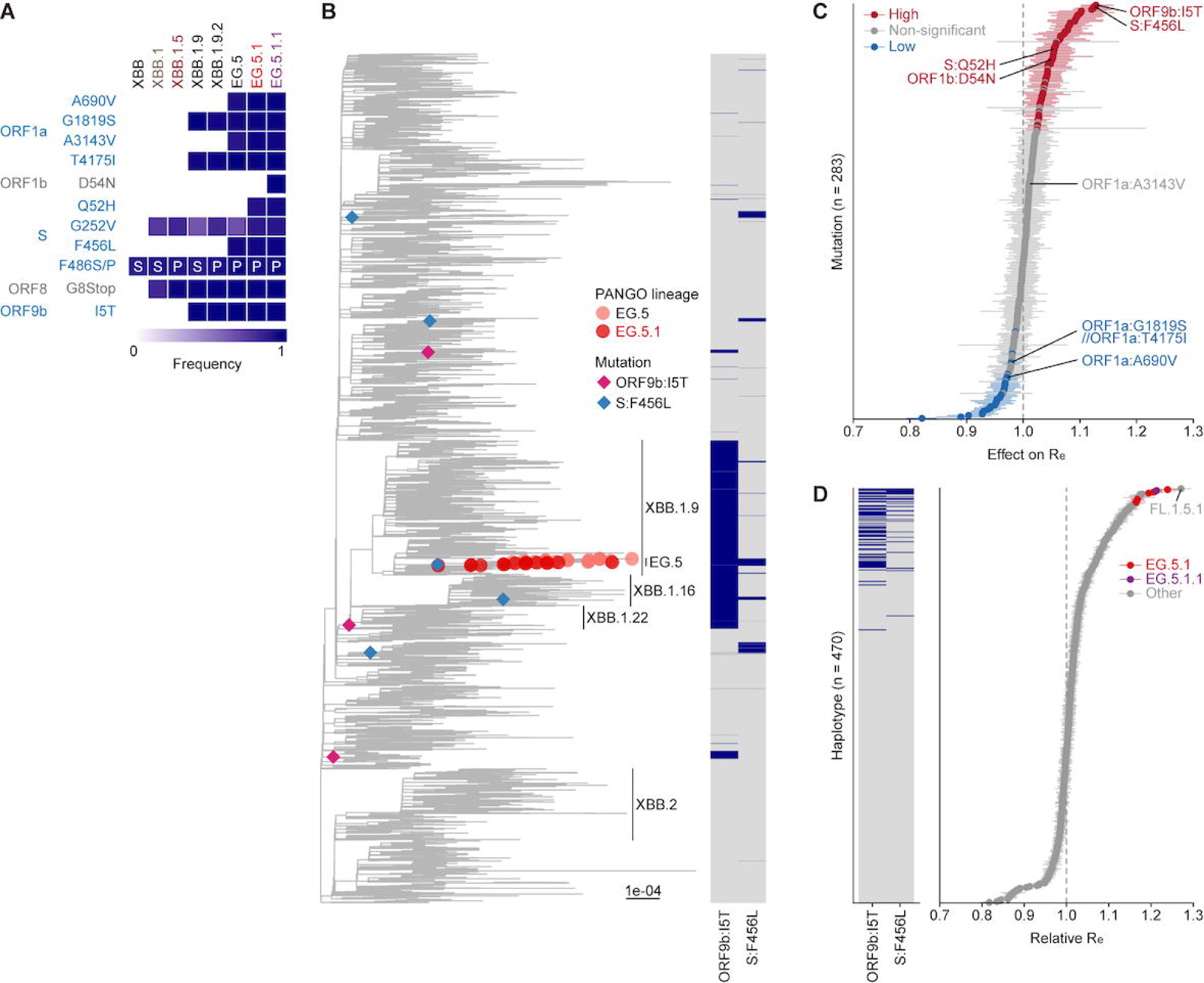
Mutations contributing to increased viral fitness of EG.5.1. (**A**) Frequency of mutations in the EG.5, EG.5.1, EG.5.1.1, and other representative XBB subvariants. Only mutations with a frequency >0.5 in at least one but not all subvariants of interest are shown. (**B**) A phylogenetic tree of SARS-CoV-2 in the XBB lineage. Only genomic sequences of SARS-CoV-2 isolates in XBB, XBB.1, EG.5, and EG.5.1 subvariants are marked with tip labels. The ultrafast bootstrap value of the common ancestor of XBB.1.9, XBB.1.16, and XBB.1.22 and that of the MRCA of EG.5 are 28 and 100, respectively. The heatmap on the right represents the presence or absence of ORF9b:I5T and S:F456L substitutions in each SARS-CoV-2 isolate. A diamond symbol represents an inferred common ancestor with an occurrence of ORF9b:I5T or S:F456L substitution. Only the occurrence events of ORF9b:I5T and S:F456L substitutions at an internal node having at least 10 descendant tips are shown. The scale bar denotes genetic distance. (**C**) Effect of substitutions in the 12 SARS-CoV-2 proteins on relative R_e_. The genome surveillance data for SARS-CoV-2 in XBB lineages circulated in the USA from December 1, 2022 to September 15, 2023 was analyzed. The posterior mean (dot) and 95% Bayesian confidential interval (CI; error bar) are shown. A group of highly co-occurred substitutions was treated as a substitution cluster. Substitutions specifically present in EG.5.1 or EG.5.1.1 compared with XBB.1.5 are labeled. (**D**) Relative R_e_ of SARS-CoV-2 haplotypes in the XBB lineage. The value for the major haplotype of XBB.1.5 is set at 1. The posterior mean (dot) and 95% Bayesian CI (error bar) are shown. The left heatmap represents the presence or absence of the ORF9b:I5T and S:F456L substitutions in each haplotype.

To test whether the convergent substitutions S:F456L and ORF9b:I5T have contributed to the increased viral fitness of EG.5.1, we performed phylogenetic and epidemic dynamics analyses using the genome surveillance data obtained from GISAID (https://gisaid.org/). First, we traced the occurrence events of ORF9b:I5T and S:F456L substitutions throughout the diversification of XBB lineages and investigate how often each substitution has occurred, which is likely to indicate the effect of substitution on viral fitness (**Figure 1B**)^5,6^. We reconstructed a phylogenetic tree of XBB lineage including 248 PANGO lineages. Subsequently, we inferred the state of presence or absence of ORF9b:I5T and S:F456L substitutions in ancestral nodes and pinpointed where each substitution occurred. We detected three and five occurrence events of ORF9b:I5T and S:F456L substitutions, respectively, supporting that these substitutions have occurred convergently during the XBB diversification (**Figure 1B**). Considering the evolutionary path of EG.5 lineage, the ORF9b:I5T substitution occurred first in a common ancestor of XBB.1.9, XBB.1.16, and XBB.1.22 lineages, which share this substitution. The S:F456L substitution occurred later in the most recent common ancestor of EG.5 lineage.

Next, we estimated the effect of ORF9b:I5T and S:F456L substitutions on viral fitness (i.e., R_e_) using a Bayesian hierarchical multinomial logistic model, established in our previous study^5^. This model can estimate the effect of an amino acid substitution on R_e_ and predict the R_e_ of a SARS-CoV-2 variant as a linear combination of the effects of individual substitutions^5^. First, we retrieved amino acid substitution profiles of SARS-CoV-2 in the XBB lineage circulated in the USA from December 1, 2022, to September 15, 2023, and classified the SARS-CoV-2 into haplotypes, groups of viruses sharing a unique substitution profile. These resulted in 470 haplotypes according to the profile of 283 substitutions in the 12 SARS-CoV-2 proteins. We then estimated the effect of each substitution on R_e_ and predicted the R_e_ of each haplotype using our model. Our modeling analysis suggests that ORF9b:I5T and S:F456L substitutions have the strongest and second-strongest positive effects on R_e_ among the substitutions we investigated, respectively (**Figure 1C, Supplementary Table 1**), whereas S:Q52H and ORF1b:D54N substitutions have a weaker positive effect on R_e_ (**Figure 1C**). Furthermore, we showed that haplotypes with ORF9b:I5T or S:F456L substitutions tend to show higher R_e_. In particular, haplotypes with both ORF9b:I5T and S:F456L substitutions, including EG.5, EG.5.1, and FL.1.5.1 (XBB.1.9.1.1.5.1), exhibit the highest R_e_ among the haplotypes we investigated (**Figure 1D, Supplementary Table 2**). FL.1.5.1 is a descendant lineage of XBB.1.9 harboring S:F456L substitution which is independent of the EG.5 lineage. Altogether, our analyses suggest that the increased viral fitness of EG.5.1 is primarily due to the ORF9b:I5T and S:F456L convergent substitutions.

### Growth kinetics of EG.5.1 and EG.5.1.1 *in vitro*

To investigate the growth kinetics of EG.5.1 and EG.5.1.1 in *in vitro* cell culture systems, we inoculated clinical isolates of Delta, XBB.1.5, EG.5.1, and EG.5.1.1 into multiple cell cultures. In Vero cells (**Figure 2A**), VeroE6/TMPRSS2 cells (**Figure 2B**) and 293-ACE2/TMPRSS2 cells (**Figure 2C**), the replication kinetics of Delta and XBB1.5 were comparable. On the other hand, the growth kinetics of EG.5.1 and EG.5.1.1 in these three cell cultures were significantly lower than that of XBB.1.5 (**Figures 2A-2C**). In Calu-3 cells (**Figure 2D**) and airway organoid-derived air-liquid interface (ALI) model (**Figure 2E**), while the replication kinetics of Delta was greater than that of XBB.1.5, those of XBB.1.5, EG.5.1, and EG.5.1.1 were comparable. In human iPSC-derived alveolar epithelial cells (**Figure 2F**), the replication kinetics of XBB.1.5 was slightly decreased compared with Delta, and EG.5.1 replication was lower than that of XBB.1.5. EG.5.1.1 showed the poorest replication capacity among the variants tested.

**Figure 2.**
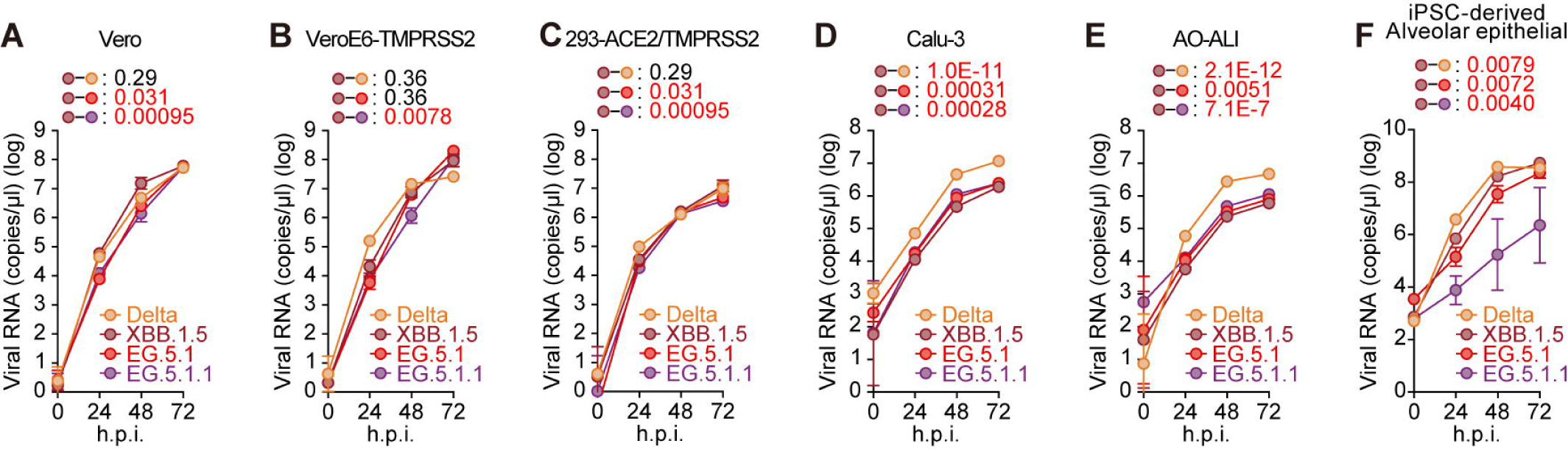
Growth kinetics of EG.5.1 and EG.5.1.1. Clinical isolates of Delta, XBB.1.5, EG.5.1, and EG.5.1.1 were inoculated into Vero cells (**A**), VeroE6/TMPRSS2 cells (**B**), 293-ACE2/TMPRSS2 cells (**C**), Calu-3 cells (**D**), airway organoids-derived ALI model (**E**), and human iPSC-derived lung alveolar cells (**F**). The copy numbers of viral RNA in the culture supernatant (**A**–**F**) were routinely quantified by RT-qPCR.

### Sensitivity of EG.5.1 and EG.5.1.1 to antiviral drugs

We then evaluated the sensitivity of EG.5.1 and EG.5.1.1 to three antiviral drugs, Remdesivir, Ensitrelvir, and Nirmatrelvir (also known as PF-07321332). Clinical isolates of Delta and XBB.1.5 were used as controls. These viruses were inoculated into human iPSC-derived lung organoids, a physiologically relevant model, and treated with three antiviral drugs. Nirmatrelvir showed the strongest antiviral effects and no differences in antiviral efficacy were observed between four variants (EC_50_ = 0.41 nM, 0.62 nM, 0.88 nM, and 0.82 nM for Delta, XBB.1.5, EG.5.1, and EG.5.1.1, respectively) (**Figure 3**). Similarly, Remdesivir and Ensitrelvir showed significant antiviral effects to these four isolates tested (**Figure 3**).

**Figure 3.**
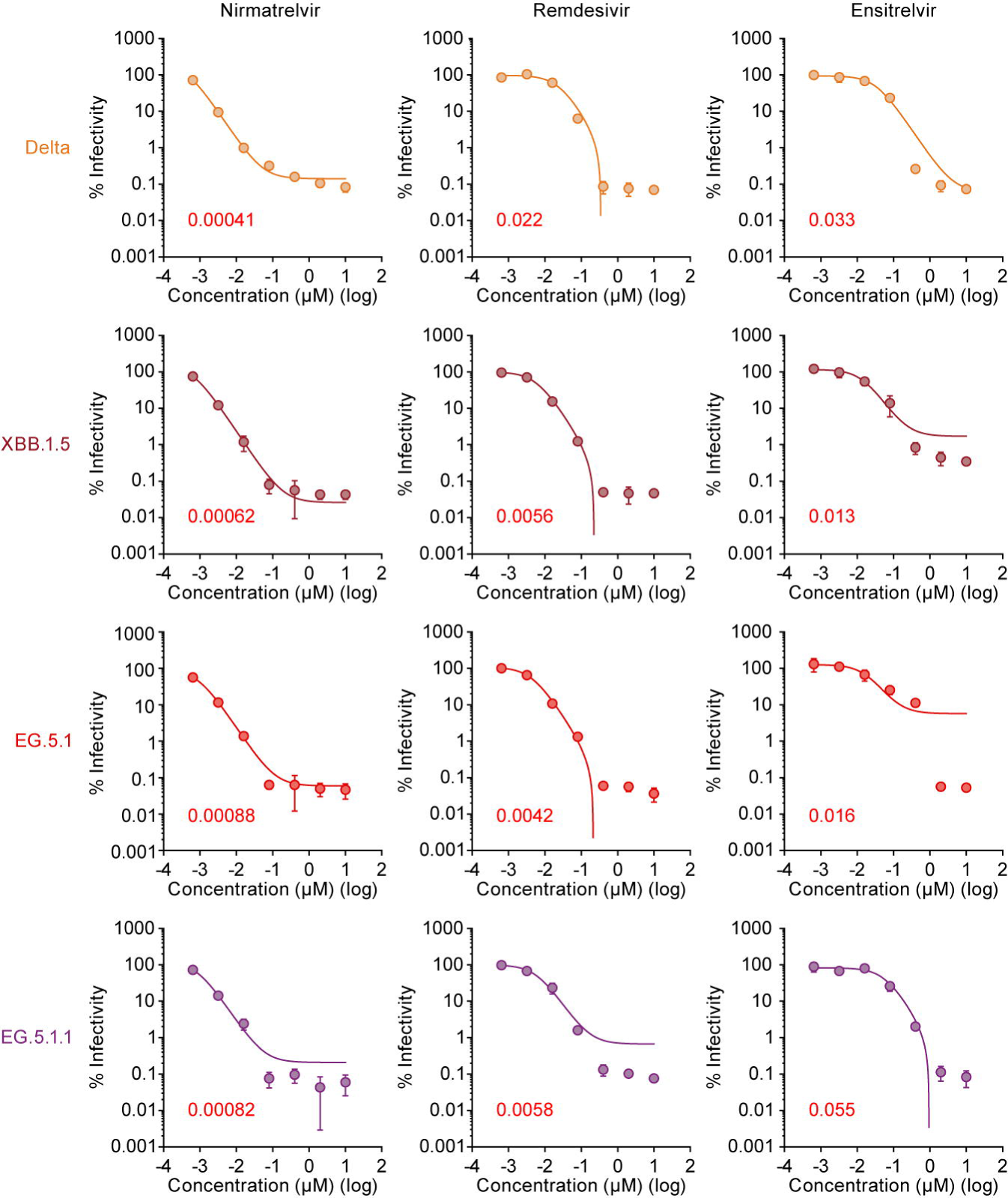
Effects of four antiviral drugs against EG.5.1 and EG.5.1.1 in human iPSC-derived lung organoids. Antiviral effects of the three drugs [Remdesivir, Ensitrelvir, and Nirmatrelvir (also known as PF-07321332)] in human iPSC-derived lung organoids. The assay of each antiviral drugs was performed in triplicate, and the 50% effective concentration (EC_50_) was calculated.

### ACE2 binding affinity of EG.5.1 S

The binding affinity of EG.5.1 S receptor binding domain (RBD) was measured by yeast surface display^7,8,10,16,19,34,36,40^. Consistent with our previous reports^2,13^, the S RBD of XBB.1.5 exhibited the lowest K_D_ value when compared to those of XBB.1 and XBB.1.16 (**Figure 4A**). Additionally, we showed that the K_D_ value of EG.5.1 S RBD was significantly higher than that of XBB.1.5 (**Figure 4A**). Similar to the observation of pseudovirus assay^4^, our data suggest that the infectious potential of EG.5.1 is not greater than that of XBB.1.5.

**Figure 4.**
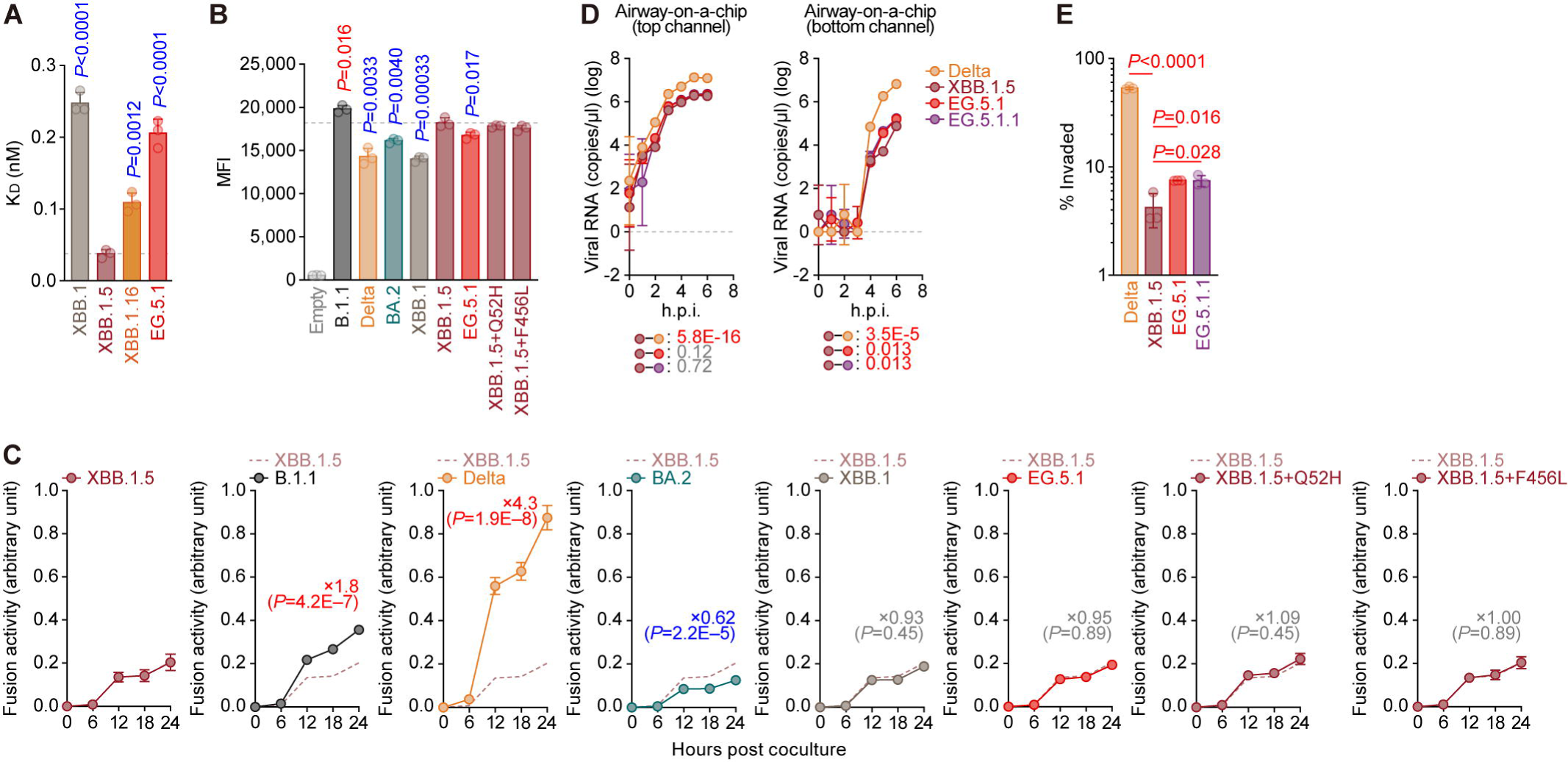
Fusogenicity of EG.5.1. **(A)** Binding affinity of the receptor binding domain (RBD) of SARS-CoV-2 spike (S) protein to angiotensin-converting enzyme 2 (ACE2) by yeast surface display. The dissociation constant (K_D_) value indicating the binding affinity of the RBD of the SARS-CoV-2 S protein to soluble ACE2 when expressed on yeast is shown. The horizontal dashed line indicates value of XBB.1.5. **(B)** Mean fluorescence intensity (MFI) of the surface S expression level in HEK293 cells. (**C**) SARS-CoV-2 S protein-mediated membrane fusion assay in Calu-3/DSP_1-7_ cells. (**D, E**) Clinical isolates of Delta, XBB.1.5, EG.5.1, and EG.5.1.1 were inoculated into an airway-on-a-chip system. The copy numbers of viral RNA in the top and bottom channels of an airway-on-a-chip were routinely quantified by RT-qPCR (**D**). The percentage of viral RNA load in the bottom channel per top channel at 6 d.p.i. (i.e., % invaded virus from the top channel to the bottom channel) is shown (**E**). Assays were performed in triplicate (**B, D, E**) or quadruplicate (**C**). The presented data are expressed as the average ± standard deviation (SD) (**B-E**). For panel **C**, statistical differences between XBB.1.5 S and each S variant across timepoints were determined by multiple regression and *P* values are indicated in each graph. The 0 h data were excluded from the analyses. The FWERs (Family-wise error rates) calculated using the Holm method are indicated in the figures.

### Fusogenicity of EG.5.1 S

The fusogenicity of EG.5.1 S protein was measured by the SARS-CoV-2 S protein-mediated membrane fusion assay^1,5,14-21^ using Calu-3/DSP_1-7_ cells. Compared to the XBB.1.5 S protein, the surface expression levels of the S proteins of Delta, BA.2, XBB.1, and EG.5.1 were reduced, while B.1.1 S protein was expressed higher on the surface of HEK293 cells (**Figure 4B**). The S:Q52H and S:F456L, hallmark amino acid substitutions of EG.5.1 S, did not affect the surface expression level of XBB.1.5 S (**Figure 4B**).

As previously reported^1,16,17,21^, the Delta S protein exhibited the greatest fusogenicity, while the BA.2 S protein exhibited the weakest fusogenicity (**Figure 4C**). Also, the XBB.1 S protein exhibited comparable fusogenicity to the XBB.1.5 S protein^22^. Here we found that the fusogenicity of EG.5.1 S was comparable to that of XBB.1.5 S, and the Q52H and F456L substitutions did not affect fusogenicity of XBB.1.5 S (**Figure 4C**). These results suggest that the EG.5.1 S protein exhibits comparable fusogenicity to XBB.1 and XBB.1.5 S proteins.

### Impact of EG.5.1 and EG.5.1.1 infection on the epithelial-endothelial barrier

To assess the effects of EG.5.1 and EG.5.1.1 infection on the airway epithelial and endothelial barriers, we employed an airway-on-a-chip system. The quantity of viruses that infiltrates from the top channel to the bottom channel reflects the capacity of viruses to breach the airway epithelial and endothelial barriers^1,5,19,22,23^. Notably, the percentage of virus that infiltrated the bottom channel of the EG.5.1- and EG.5.1.1-infected airway-on-a-chip was comparable to that of the XBB.1.5-infected airway-on-a-chip (**Figures 4D and 4E**). Together with the findings of the S-based fusion assay (**Figure 4C**), these results suggest that the fusogenicity of EG.5.1 and EG.5.1.1 is comparable to that of XBB.1.5.

### Structural characteristics of EG.5.1 S protein

To gain structural insights into EG.5.1 S protein, the structures of the EG.5.1 S ectodomain alone were determined by cryoelectron microscopy (cryo-EM) analysis. The EG.5.1 S ectodomain was reconstructed as two closed states and a 1-up state at resolutions of 2.50 Å, 2.89 Å and 3.34 Å, respectively (**Figure 5A, Supplementary Figures 1A-B, and Supplementary Table 3**). The two closed states in EG.5.1 show structural differences in the orientation of the RBD and the loop structure at the protomer interface (**Figure 5A** and **Supplementary Figure 1C)**, as observed in XBB.1 and XBB.1.5^1^, therefore, these two closed states were defined as closed-1 and closed-2, respectively. In addition, a 1-up state was also observed in EG.5.1, which could not be observed in XBB.1 and XBB.1.5. XBB variant was derived from recombination of BJ.1.1 with BM.1.1.1, a descendant of BA.2.75^1^, two closed and a 1-up states are observed in BA.2.75 S like EG.5.1 S^19,24^. Thus, from BA.2.75 through XBB to EG.5.1, there exist conformational differences among the representative structures of the spike protein of these variants. To examine the reason for transition of spike protein conformation, we compared the structures of XBB.1.5 and EG.5.1^22^. While closed-1 state of XBB.1.5 and EG.5.1 share a nearly identical overall structure, the relative orientation of RBD in the closed-2 state show a minor displacement (**Figure 5B**). EG.5.1 has the Q52H substitution in the NTD and the F456L substitution in the RBD compared to XBB.1.5 (**Supplementary Figures 1D-E**), especially the F456L substitution is located at the interface between protomers in the closed-2 state (**Figure 5C**). When focusing on the interactions of F456L, in XBB.1.5 S, F456 was located at a distance of 3.8 Å from P373, whereas in EG.5.1, F456L and P373 exhibited a distance of 10.1 Å, thus resolving hydrophobic interactions (**Figure 5C**). This structural difference suggests that the closed-2 state of EG.5.1 exhibits a weaker RBD packing compared to XBB.1/XBB.1.5, allowing for EG.5.1 S to more likely transit to the 1-up state. It has been previously reported that the F486 residue stabilizes the 1-up conformation by interacting to the up RBD^25,26^. To verify the position of amino-acid residue 486 in RBD and up RBDs of EG.5.1 S, we focused on the interface between down and up RBDs in the 1-up state (**Supplementary Figure 1A**). Although the details of the interaction are unclear due to resolution limitations, the P486 residue was found to be in contact with the up RBD residue, suggesting that it may also contribute to the stabilization of the 1-up state (**Figure 5D**).

**Figure 5.**
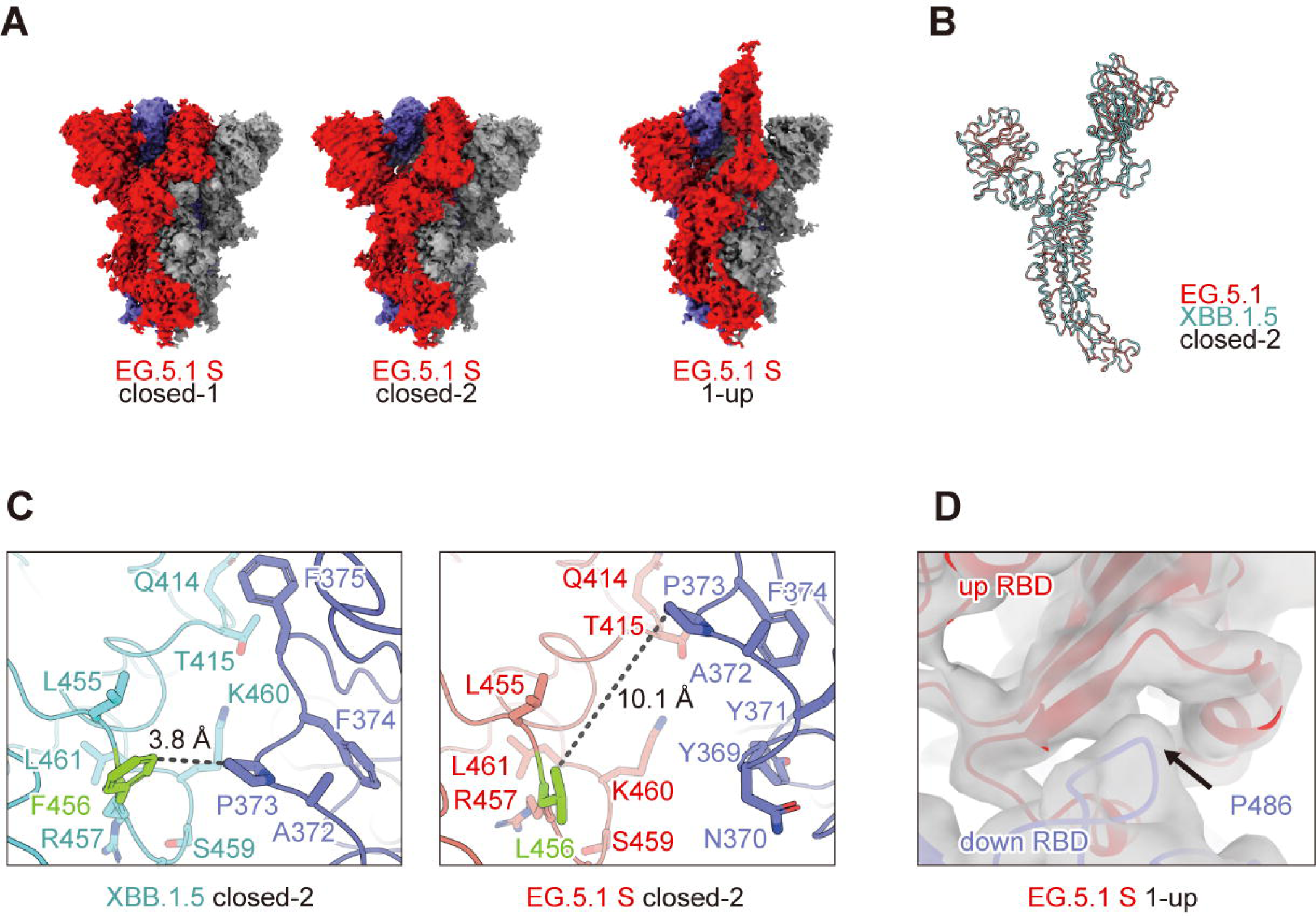
Overall cryo-EM maps and structures of SARS-CoV-2 EG.5.1 S protein. (**A**) Cryo-EM maps of EG.5.1 S protein trimer closed-1 state (**Left**), closed-2 state (**Middle**) and 1-up state (**Right**). Each protomer is colored red, blue, gray. (**B**) Superimposed structures of EG.5.1 (red) and XBB.1.5 (cyan) S protomers in closed-2 state. The models were superposed on the Cα atoms of the corresponding residues in the S2 region (RMSD = 0.244). (**C**) Close-up views of the amino-acid residues 456 and 373 in closed-2 structures. (**Left**) F456 at the protomer interface in the XBB.1.5 S RBD region makes hydrophobic contact with P373 in adjacent protomer at a distance of 3.8 Å. (**Right**) F456L substitution causes loss of hydrophobic contact with P373, up to 10.1 Å away. (**D)**. Close-up view of the interface between up RBD and adjacent down protomer. The model of EG.5.1 closed-2 RBD was rigid-fitted to the corresponding region of the cryo-EM map of EG.5.1 S protein 1-up state.

### Virological characteristics of EG.5.1 and EG.5.1.1 *in vivo*

To investigate the virological features of EG.5.1 and its variant EG.5.1.1 *in vivo*, clinical isolates of Delta, XBB.1.5, EG.5.1, and EG.5.1.1 (10,000 TCID_50_) were intranasally inoculated into hamsters under anesthesia. Consistent with our previous studies^1,5,15,16,19^, Delta infection resulted in weight loss (**Figure 6A, left**). The body weights of the hamsters infected with XBB.1.5, EG.5.1 and EG.5.1.1 were comparable and significantly lower than that of uninfected hamsters (**Figure 6A, left**).

**Figure 6.**
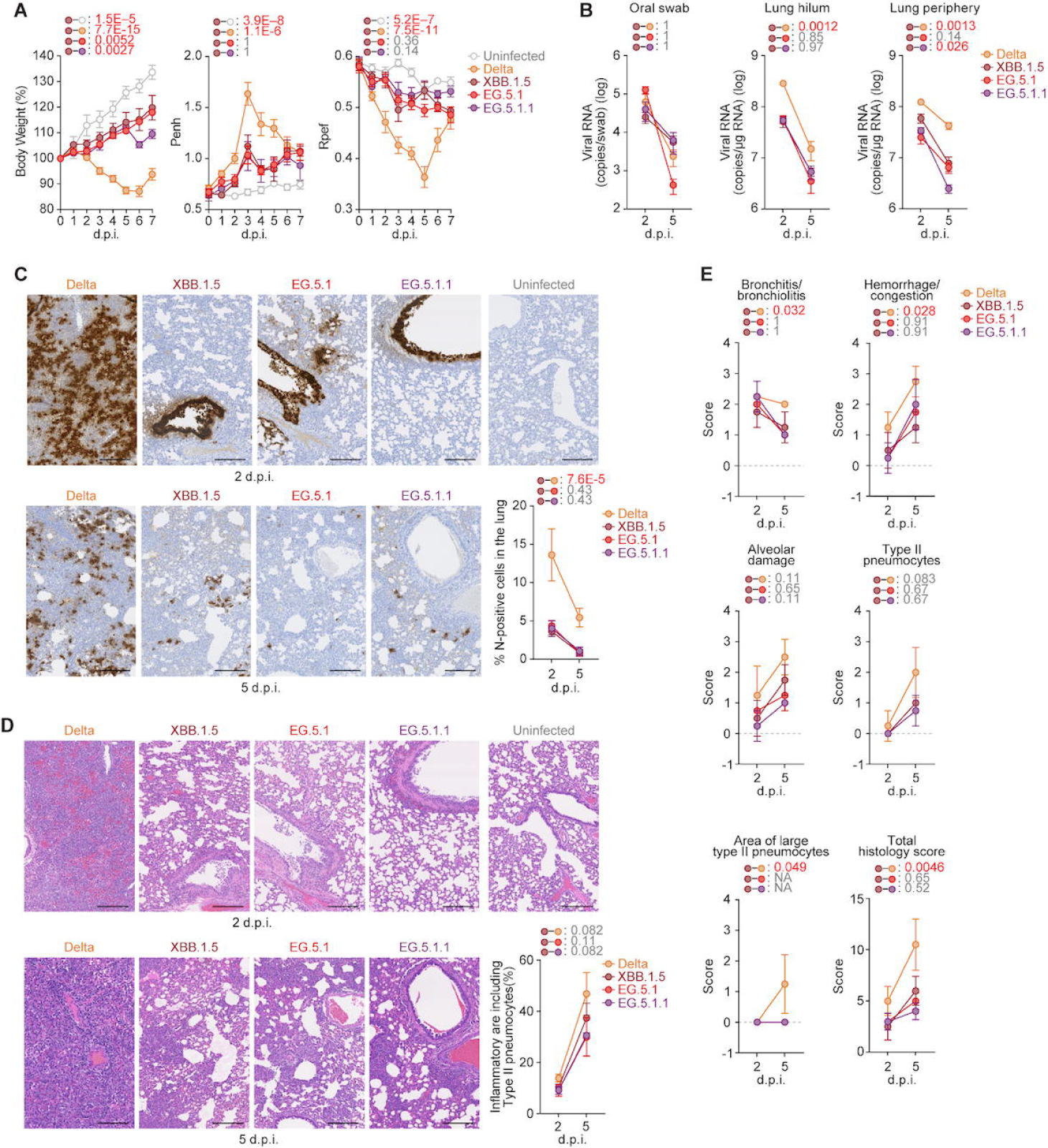
Virological characteristics of EG.5.1 and EG.5.1.1 *in vivo*. Syrian hamsters were intranasally inoculated with EG.5.1, EG.5.1.1, XBB.1.5, and Delta. Six hamsters of the same age were intranasally inoculated with saline (uninfected). Six hamsters per group were used to routinely measure the respective parameters (**A**). Four hamsters per group were euthanized at 2 and 5 d.p.i. and used for virological and pathological analysis (**C–E**). (**A**) Body weight, Penh, and Rpef values of infected hamsters (*n* = 6 per infection group). (**B**) (**Left**) Viral RNA loads in the oral swab (*n* = 6 per infection group). (**Middle** and **right**) Viral RNA loads in the lung hilum (**middle**) and lung periphery (**right**) of infected hamsters (*n* = 4 per infection group). (**C**) IHC of the viral N protein in the lungs at 2 d.p.i. (**left**) and 5 d.p.i. (**right**) of infected hamsters. Representative figures (N-positive cells are shown in brown). (**D**) H&E staining of the lungs of infected hamsters. Representative figures are shown in (**D**). Uninfected lung alveolar space is also shown. The raw data are shown in **Supplementary Figure 1**. (**E**) Histopathological scoring of lung lesions (*n* = 4 per infection group). Representative pathological features are reported in our previous studies^15-17,32-35^. In **A–C, E**, data are presented as the average ± SEM. In **C**, each dot indicates the result of an individual hamster. In **A,B**,**E**, statistically significant differences between EG.5.1, EG5.1.1 and other variants across timepoints were determined by multiple regression. In **B**,**E**, the 0 d.p.i. data were excluded from the analyses. The FWERs calculated using the Holm method are indicated in the figures. In **C**, the statistically significant differences between EG.5.1, EG.5.1.1 and other variants were determined by a two-sided Mann–Whitney *U* test. In **C** and **D**, each panel shows a representative result from an individual infected hamster. Scale bars, 200 μm (**C, D**).

We then analyzed the pulmonary function of infected hamsters as reflected by two parameters, enhanced pause (Penh) and the ratio of time to peak expiratory flow relative to the total expiratory time (Rpef). Delta infection resulted in significant differences in these two respiratory parameters compared to XBB.1.5 infection (**Figure 6A, middle and right**), suggesting that Delta is more pathogenic than XBB.1.5. On the other hand, the Penh and Rpef values of EG.5.1-, EG.5.1.1-, and XBB.1.5-infected hamsters were comparable (**Figure 6A, middle and right**), suggesting that the pathogenicity of EG.5.1 variants is similar to that of XBB.1.5 in hamsters.

To evaluate the viral spread in infected hamsters, we routinely measured the viral RNA load in the oral swab. Although the viral RNA load of EG.5.1-infected hamsters were significantly higher than that of XBB.1.5-infected hamster at 2 d.p.i., the EG.5.1 RNA load was acutely decreased and was significantly lower than the XBB.1.5 RNA load at 5 d.p.i (**Figure 6B, left**).

We then compared the viral spread in the respiratory tissues. We collected the lungs of infected hamsters at 2 and 5 d.p.i., and the collected tissues were separated into the hilum and periphery regions. The viral RNA loads in both the lung hilum and periphery regions of Delta-infected hamsters were significantly higher than those of the other three Omicron subvariants. The viral RNA loads in both lung regions of EG.5.1-infected hamsters were slightly lower than those of XBB.1.5-infected hamsters (**Figure 6B, middle and right**). In the lung hilum region, the viral RNA load of EG.5.1.1-infected hamsters were comparable to that of XBB.1.5-infected hamsters at 2 and 5 d.p.i. (**Figure 6B, middle**). However, in the lung periphery region, the EG.5.1.1 RNA load was significantly lower than the XBB.1.5 RNA load (**Figure 6B, right**). These results suggest that the viral spreading efficacy of EG.5.1 and EG.5.1.1 in the lung is lower than that of XBB.1.5.

To further investigate of the viral spread in the respiratory tissues of infected hamsters, we analyzed the formalin-fixed right lungs of infected hamsters at 2 and 5 d.p.i. by carefully identifying the four lobules and lobar bronchi sectioning each lobe along with the bronchial branches and performed immunohistochemical (IHC) analysis targeting viral nucleocapsid (N) protein. Consistent with our previous studies^1,5,15-19,22^, at 2 d.p.i., the N-positive cells were strongly detected in Delta-infected hamsters in the alveolar space around the bronchi/bronchioles (**Figure 6C**). In the three Omicron subvariants, the percentage of N-positive cells in the lungs of EG.5.1- and EG.5.1.1-infected hamsters were comparable to that of XBB.1.5-infected hamsters (**Figure 6C and Supplementary Figure 2A**). At 5 d.p.i., N-positive cells were detected in the peripheral alveolar space in Delta-infected hamsters, while the N-positive areas of EG.5.1-, EG.5.1.1- and XBB.1.5-infected hamsters were slightly detectable in the peripheral alveolar space (**Figure 6C and Supplementary Figure 2B**). There was no significant difference in the N-positive area of three Omicron subvariants (**Supplementary Figures 2A and 2B**).

### Intrinsic pathogenicity of EG.5.1 and EG.5.1.1

To investigate the intrinsic pathogenicity of EG.5.1 and EG.5.1.1, histopathological analyses were performed according to the criteria described in our previous study^16^. At 2 d.p.i., inflammation was limited in bronchi/bronchioles in the hamsters infected with EG.5.1, EG.5.1.1 and XBB.1.5 (**Figure 6D and Supplementary Figure 3A**). On the other hand, alveolar damage around the bronchi was prominent in Delta-infected hamsters (**Figure 6D**). At 5 d.p.i., although the alveolar architecture was totally destroyed by the alveolar damage or the expansion of type II pneumocytes in Delta-infected hamsters, alveolar architecture was preserved in the three Omicron subvariant-infected hamsters (**Figures 6D, 6E and Supplementary Figure 3B**). Consistent with our previous studies^1,5,15-19,22^, all five histological parameters and the total histology score of the Delta-infected hamsters were greatest (**Figure 6E**). On the other hand, these scores were comparable in the three Omicron subvariant-infected hamsters (**Figures 6D and 6E**).

### Impact of ORF9b:I5T on IFN-I inhibition and viral growth kinetics

As shown in **Figure 1**, our epidemic dynamics analyses suggested that ORF9b:I5T substitution contributes to the increased viral fitness in XBB lineages. In addition to ORF9b:I5T, EG.5.1 has a substitution and a deletion in this protein compared with Wuhan-Hu-1 (WH1)^27^, both of which are conserved across Omicron lineages (**Figure 7A**). Since previous studies demonstrated that ORF9b inhibits IFN-I signaling^7-10^, we hypothesized that the I5T substitution affects the anti-IFN-I function of ORF9b. To address this possibility, we used the expression plasmid for the ORF9b protein of WH1^27^ and compared its anti-IFN-I activity with that of ORF6, another viral anti-IFN-I antagonist, which we showed previously^28^. As shown in **Figure 7B**, the anti-IFN-I activity of WH1 ORF9b was less than that of WH1 ORF6. We then assessed the anti-IFN-I activity of ORF9b of some SARS-CoV-2 Omicron subvariants such as XBB.1.5, XBB.1.16, and EG.5.1. Although some values of Omicron subvariants were different from that of WH1 with statistically significance, the anti-IFN-I activity was not clearly different (**Figure 7C**). These findings suggest that the I5T substitution does not critically affect the anti-IFN-I activity of ORF9b.

**Figure 7.**
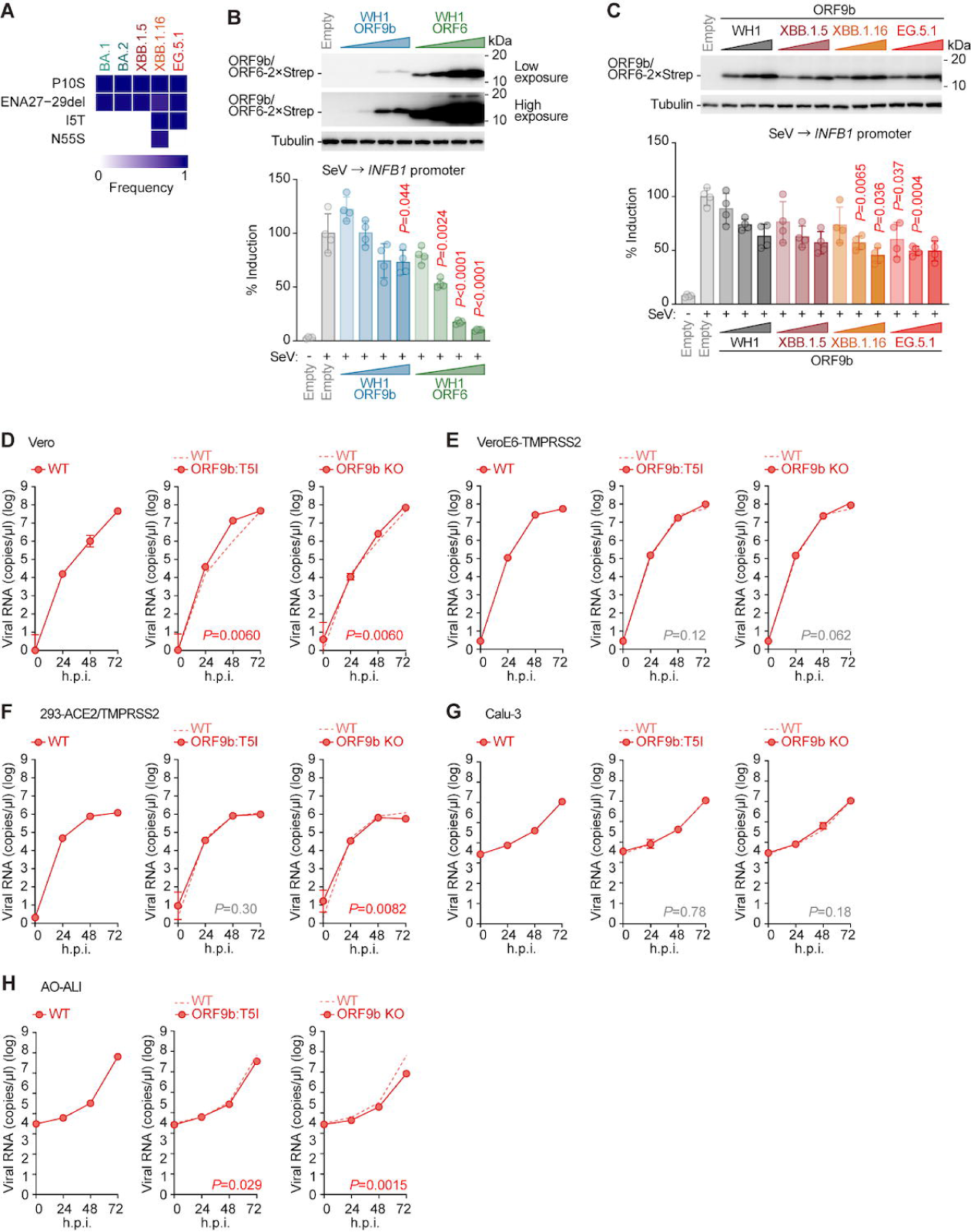
Impact of the I5T substitution of ORF9b on innate immune response and viral growth. (**A–C**) Anti-IFN-I effect of ORF9b:I5T. (**A**) Frequency of mutations in ORF9b of BA.1, BA.2, XBB.1.5, XBB.1.16, and EG.5.1. Only mutations with a frequency >0.5 in at least a subvariant are shown. (**B**) Comparison of the anti-IFN-I effect and expression levels between ORF9b and ORF6 in HEK293 cells. HEK293 cells were cotransfected with plasmids expressing 2×Strep-tagged ORF9b or ORF6 and p125Luc. 24 h after transfection, cells were infected with SeV (MOI 100). 24 h after infection, cells were harvested for western blotting (**top**) and a luciferase assay (**bottom**). (**C**) Comparison of the anti-IFN-I effect and expression levels of ORF9b among SARS-CoV-2 variants in HEK293 cells. HEK293 cells were cotransfected with plasmids expressing 2×Strep-tagged ORF9b variants and p125Luc. 24 h after transfection, cells were infected with SeV (MOI 100). 24 h after infection, cells were harvested for western blotting (**top**) and a luciferase assay (**bottom**). For Western blotting (**B, top** and **C, top**), the input of cell lysate was normalized to TUBA, and one representative result out of three independent experiments is shown. kDa, kilodalton. For the luciferase assay (**B, bottom** and **C, bottom**), the value was normalized to the unstimulated, empty vector-transfected cells (no SeV infection). (**D–H**) Three recombinant SARS-CoV-2, rEG.5.1 WT, rEG.5.1 ORF9b:T5I (ORF9b:T5I), and rEG.5.1 ORF9b KO (ORF9b KO) were inoculated into Vero cells (**D**), VeroE6/TMPRSS2 cells (**E**), 293-ACE2/TMPRSS2 cells (**F**), Calu-3 cells (**G**), and airway organoids-derived ALI model (**H**). The copy numbers of viral RNA in the culture supernatant (**D–H**) were routinely quantified by RT-qPCR. The dashed red line indicates the results of WT. In **B**, the statistically significant differences between the stimulated, empty vector-transfected cells (SeV infection) and the stimulated, ORF9b or ORF6 expression vector-transfected cells were determined by a two-sided Student’s *t* test. In **C**, the statistically significant differences between the stimulated, ORF9b expression vector-transfected cells and the stimulated, ORF6 expression vector-transfected cells at the same dose respectively were determined by a two-sided Student’s *t* test.

To further evaluate the impact of ORF9b:I5T on viral growth kinetics in *in vitro* cell culture systems, we prepared three recombinant SARS-CoV-2, rEG.5.1 [wildtype (WT)], rEG.5.1 ORF9b:T5I (ORF9b:T5I), and rEG.5.1 ORF9b KO (ORF9b KO) by reverse genetics and inoculated it into multiple cell cultures. As shown in **Figure 7D-H**, viral growth kinetics of ORF9b:T5I was comparable to that of WT in all tested cell cultures, suggesting that the ORF9b:I5T does not affect the viral replication efficacy. Similarly, the growth kinetics of WT and ORF9b KO were comparable (**Figure 7D-H**). These findings suggest that ORF9b does not have a crucial role on viral replication at least in *in vitro* cell culture systems.

## Discussion

In this study, we performed phylogenetic and epidemic dynamics modeling analyses using viral sequence data and showed the data suggesting that two hallmark mutations in the EG.5.1 lineage, S:F456L and ORF9b:I5T, are critical to the increased viral fitness (i.e., R_e_). We then experimentally addressed the growth kinetics, sensitivity to clinically available antiviral compounds, fusogenicity and pathogenicity of EG.5.1 and EG.5.1.1 variants. Our experimental results suggest that the virological features of EG.5.1 and EG.5.1.1 are almost comparable to that of XBB.1.5. We further reveal the structure of EG.5.1 S by cryo-EM and describe the structural difference between the EG.5.1 S and XBB.1.5 S. Moreover, we assessed the impact of ORF9b:I5T on the function of IFN-I antagonism by ORF9b, while its impact was negligible at least in our experimental setup. Furthermore, the experiments using the recombinant EG.5.1 viruses by reverse genetics showed that the impact of ORF9b:I5T on viral growth is not observed.

We have shown the evidence suggesting that the fusogenicity of S protein in *in vitro* cell cultures (particularly in Calu-3 cells) is closely associated with viral intrinsic pathogenicity in hamsters^15,16,18^. Consistent with our assumption, we demonstrated the fusogenicity of EG.5.1 S is comparable to that of XBB.1.5 S (**Figure 4C**), and the infection experiment using airway-on-a-chip showed that the potential of EG.5.1 to invade epithelial-endothelial barrier, which reflects viral fusogenicity, is similar to that of XBB.1.5 (**Figures 4D and 4E**). Moreover, in the experiments using hamsters, the intrinsic pathogenicity of EG.5.1 is also indistinguishable to that of XBB.1.5 (**Figure 6**). Our results suggest that the viral virulence is not modulated by the mutations accumulated in the EG.5.1 genome when compared to XBB.1.5.

The ACE2 binding assay *in vitro* showed that the K_D_ value of EG.5.1 S RBD is significantly higher than that of XBB.1.5 S RBD (**Figure 4A**). Consistent with this observation, we have previously found that the pseudovirus infectivity of EG.5.1 is also lower than that of XBB.1.5^4^. Additionally, the growth kinetics of EG.5.1 does not outweigh that of XBB.1.5, while its extent is dependent on the cell types used (**Figure 2**). Moreover, in hamsters, the spreading efficiency of EG.5.1 is comparable to XBB.1.5 (**Figures 6B and 6C**). These observations suggest that the growth capacity of EG.5.1 is similar to that of XBB.1.5. On the other hand, as explained in the Introduction, recent studies including ours^4,11^ showed that EG.5.1 exhibits significantly greater immune resistance to the humoral immunity induced by XBB breakthrough infection than XBB.1.5, and the S:F456L substitution is responsible for this immunological phenotype. Altogether, these observations suggest that the epidemic spread of the EG.5.1 lineage by outcompeting an S:F486P-bearing XBB subvariants including XBB.1.5, is not due to increased viral growth capacity, but rather to increased immune evasion capacity from the humoral immunity induced by XBB breakthrough infection.

The structure of EG.5.1 S alone revealed in this study provides an opportunity to discuss the impact of the substitutions in S proteins occurring in variants since BA.2.75 on a series of conformational changes from BA.2.75 through XBB.1 and XBB.1.5 to EG.5.1^1,19,22^. It has been reported that the F486 residue stabilizes the 1-up state by hydrophobic interactions with the up RBD in the 1-up conformation^25,26^. In XBB.1 S, the 1-up conformation is not optimal due to F486S substitution, a less bulky and hydrophilic residue. On the other hand, XBB.1.5 and EG.5.1 S acquired commonly the F486P substitution, these variants are thought to have the potential to stabilize an up RBD conformation, but only EG.5.1 S was reconstructed in the 1-up conformation. This difference between XBB.1.5 and EG.5.1 is probably related to the F456L substitution, which resolved the interaction between protomers in closed 2 in EG.5.1 and facilitated the transition from the closed state to the 1-up one. The amino-acid substitution-dependent conformational transitions illuminated by this study provide an understanding of metastable pre-fusions state of the spike protein in omicron subvariant, BA.2.75, XBB.1, XBB.1.5 and EG.5.1.

### Limitation of the study

Our epidemic dynamics modeling analysis suggests that S:F456L and ORF9b:I5T enhance viral fitness in XBB lineages (**Figure. 1**). S:F456L likely boosts viral fitness by improving the ability to escape humoral immunity induced by vaccination and natural infection^4^. On the contrary, we failed to uncover any notable effects of ORF9b:I5T on the viral properties we examined, including viral replication in cell lines or airway organoids, or the inhibition of the IFN pathway (**Figure. 7**). This discrepancy regarding the ORF9b:I5T might be attributed to two potential explanations. First, the observed effect of ORF9b:I5T on viral fitness could be a false positive. However, this seems less likely since the positive effect of ORF9b:I5T on viral fitness was supported by two independent methods: our approach and a method developed by Bloom et al., which infers the fitness effect of a mutation based on its convergent acquisition level^6^. The second explanation is that ORF9b:I5T might affect viral properties related to fitness, which we did not investigate experimentally. Indeed, our understanding of which properties of the virus are closely related to viral fitness is currently limited. For a deeper understanding of the mechanism by which the virus boosts its transmission potential, characterizing mutations in non-S proteins including ORF9b:I5T would be crucial.

In sum, our multiscale investigation revealed the virological characteristics of a most recently spreading SARS-CoV-2 variant, EG.5.1, particularly focusing on the effects of hallmark substitutions in the S (F456L) and non-S (ORF9b:I5T) proteins. As we demonstrated on a variety of SARS-CoV-2 Omicron subvariants in the past^1,2,4,5,13,15,17-19,21,22,29-31^, elucidating the virological features of newly emerging SARS-CoV-2 variants is important to consider the potential risk to human society and to understand the evolutionary scenario of the emerging virus in the human population. In particular, accumulating the knowledge of the evolution trait of newly emerging pathogenic viruses in the human population will be beneficial for future outbreak/pandemic preparedness.

## Author Contributions

Sayaka Deguchi, MST Monira Begum, Hesham Nasser, Michael Jonathan, Terumasa Ikeda, and Kazuo Takayama performed cell culture experiments.

Shuhei Tsujino and Tomokazu Tamura generated recombinant viruses.

Shuhei Tsujino, Keita Mizuma, Naganori Nao, Isshu Kojima, Tomoya Tsubo, Jingshu Li, Kumiko Yoshimatsu, Rigel Suzuki, Tomokazu Tamura, Keita Matsuno performed animal experiments.

Lei Wang, Yoshitaka Oda, Masumi Tsuda, Shinya Tanaka performed histopathological analysis. performed yeast surface display assay.

Sayaka Deguchi, Kazuo Takayama prepared human iPSC-derived lung organoids, AO-ALI, and airway-on-a-chip systems.

Sayaka Deguchi, Kazuo Takayama performed antiviral drug tests

Yuki Yamamoto and Tetsuharu Nagamoto performed generation and provision of human iPSC-derived airway and alveolar epithelial cells.

Hisano Yajima, Kaori Sasaki-Tabata and Takao Hashiguchi prepared the EG.5.1 S protein.

Tomo Nomai, Yuki Anraku, Shunsuke Kita, Katsumi Maenaka and Takao Hashiguchi determined the structure of EG.5.1 S protein.

Hiroyuki Asakura, Mami Nagashima, Kenji Sadamasu and Kazuhisa Yoshimura performed viral genome sequencing analysis.

Yasufumi Matsumura, Miki Nagao collected swab samples from COVID-19 and performed viral genome sequencing analysis.

Arnon Plianchaisuk performed bioinformatics analyses.

Jumpei Ito designed bioinformatics analyses and interpreted the results. Terumasa Ikeda, Takasuke Fukuhara, and Kei Sato designed the experiments and interpreted the results.

Jumpei Ito, Terumasa Ikeda, Kazuo Takayama, Jiri Zahradnik, Takasuke Fukuhara and Kei Sato wrote the original manuscript.

All authors reviewed and proofread the manuscript.

The Genotype to Phenotype Japan (G2P-Japan) Consortium contributed to the project administration.

## Acknowledgments

We would like to thank all members belonging to The Genotype to Phenotype Japan (G2P-Japan) Consortium. We thank Dr. Kenzo Tokunaga (National Institute for Infectious Diseases, Japan) and Dr. Jin Gohda (The University of Tokyo, Japan) for providing reagents. We thank to all members belonging to Japanese Consortium on Structural Virology (JX-Vir). We appreciate the technical assistance from The Research Support Center, Research Center for Human Disease Modeling, Kyushu University Graduate School of Medical Sciences. We gratefully acknowledge all data contributors, i.e. the Authors and their Originating laboratories responsible for obtaining the specimens, and their Submitting laboratories for generating the genetic sequence and metadata and sharing via the GISAID Initiative, on which this research is based. The super-computing resource was provided by Human Genome Center at The University of Tokyo.

This study was supported in part by AMED SCARDA Japan Initiative for World-leading Vaccine Research and Development Centers “UTOPIA” (JP223fa627001, to Kei Sato), AMED SCARDA Program on R&D of new generation vaccine including new modality application (JP223fa727002, to Kei Sato); AMED SCARDA Kyoto University Immunomonitoring Center (KIC) (JP223fa627009, to Takao Hashiguchi); AMED SCARDA Hokkaido University Institute for Vaccine Research and Development (HU-IVReD) (JP223fa627005, to Katsumi Maenaka); AMED Research Program on Emerging and Re-emerging Infectious Diseases (JP21fk0108574, to Hesham Nasser; JP21fk0108493, to Takasuke Fukuhara; JP22fk0108617 to Takasuke Fukuhara; JP22fk0108146, to Kei Sato; JP21fk0108494 to G2P-Japan Consortium, Keita Matsuno, Shinya Tanaka, Terumasa Ikeda, Takasuke Fukuhara, and Kei Sato; JP21fk0108425, to Kazuo Takayama and Kei Sato; JP21fk0108432, to Kazuo Takayama, Takasuke Fukuhara and Kei Sato; JP22fk0108534, to Takashi Irie, Terumasa Ikeda, and Kei Sato; JP22fk0108511, to Yuki Yamamoto, Terumasa Ikeda, Keita Matsuno, Shinya Tanaka, Kazuo Takayama, Takao Hashiguchi, Takasuke Fukuhara, and Kei Sato); AMED Research Program on HIV/AIDS (JP22fk0410055, to Terumasa Ikeda; and JP22fk0410039, to Kei Sato); AMED Japan Program for Infectious Diseases Research and Infrastructure (JP22wm0125008 to Keita Matsuno); AMED CREST (JP21gm1610005, to Kazuo Takayama; JP22gm1610008, to Takasuke Fukuhara; JP22gm1810004, to Katsumi Maenaka); JST PRESTO (JPMJPR22R1, to Jumpei Ito); JST CREST (JPMJCR20H4, to Kei Sato; JPMJCR20H8, to Takao Hashiguchi); JSPS KAKENHI Grant-in-Aid for Scientific Research C (22K07103, to Terumasa Ikeda); JSPS KAKENHI Grant-in-Aid for Scientific Research B (21H02736, to Takasuke Fukuhara); JSPS KAKENHI Grant-in-Aid for Early-Career Scientists (22K16375, to Hesham Nasser; 20K15767, Jumpei Ito); JSPS KAKENHI grant JP20H05873 (to Katsumi Maenaka); JSPS Core-to-Core Program (A. Advanced Research Networks) (JPJSCCA20190008, to Kei Sato); JSPS Research Fellow DC2 (22J11578, to Keiya Uriu); JSPS Leading Initiative for Excellent Young Researchers (LEADER) (to Terumasa Ikeda); World-leading Innovative and Smart Education (WISE) Program 1801 from the Ministry of Education, Culture, Sports, Science and Technology (MEXT) (to Naganori Nao); Research Support Project for Life Science and Drug Discovery [Basis for Supporting Innovative Drug Discovery and Life Science Research (BINDS)] from AMED under the Grant JP22ama121001 (to Takao Hashiguchi) and JP22ama121037 (to Katsumi Maenaka); The Cooperative Research Program (Joint Usage/Research Center program) of Institute for Life and Medical Sciences, Kyoto University (to Kei Sato and Katsumi Maenaka); International Joint Research Project of the Institute of Medical Science, the University of Tokyo (to Terumasa Ikeda, Jiri Zahradnik, and Takasuke Fukuhara); The Tokyo Biochemical Research Foundation (to Kei Sato); Takeda Science Foundation (to Terumasa Ikeda and Katsumi Maenaka); Mochida Memorial Foundation for Medical and Pharmaceutical Research (to Terumasa Ikeda); The Naito Foundation (to Terumasa Ikeda); Mitsubishi Foundation (to Kei Sato); and the project of National Institute of Virology and Bacteriology, Programme EXCELES, funded by the European Union, Next Generation EU (LX22NPO5103, to Jiri Zahradnik).

## Consortia

Hirofumi Sawa^11^, Tomoya Tsubo^11^, Zannatul Ferdous^7^, Kenji Shishido^7^, Saori Suzuki^1^, Hayato Ito^1^, Yu Kaku^6^, Naoko Misawa^6^, Kaoru Usui^6^, Wilaiporn Saikruang^6^, Yusuke Kosugi^6^, Shigeru Fujita^6^, Jarel Elgin M. Tolentino^6^, Luo Chen^6^, Lin Pan^6^, Mai Suganami^6^, Mika Chiba^6^, Ryo Yoshimura^6^, Kyoko Yasuda^6^, Keiko Iida^6^, Adam P. Strange^6^, Naomi Ohsumi^6^, Shiho Tanaka^6^, Kaho Okumura^6^, Daniel Sauter^6,36^, Isao Yoshida^20^, So Nakagawa^36^, Kotaro Shirakawa^37^, Akifumi Takaori-Kondo^37^, Kayoko Nagata^37^, Ryosuke Nomura^37^, Yoshihito Horisawa^37^, Yusuke Tashiro^37^, Yugo Kawai^37^, Rina Hashimoto^2^, Yukio Watanabe^2^, Yoshitaka Nakata^2^, Hiroki Futatsusako^2^, Ayaka Sakamoto^2^, Naoko Yasuhara^2^, Tateki Suzuki^16^, Kanako Terakado Kimura^16^, Jiei Sasaki^16^, Yukari Nakajima^16^, Ryoko Kawabata^29^, Ryo Shimizu^9^, Yuka Mugita^9^, Sharee Leong^9^, Otowa Takahashi^9^, Kimiko Ichihara^9^, Chihiro Motozono^38^, Mako Toyoda^38^, Takamasa Ueno^38^, Akatsuki Saito^39^, Maya Shofa^39^, Yuki Shibatani^39^, Tomoko Nishiuchi^39^, Prokopios Andrikopoulos^4^, Aditi Konar^4^

^36^Tokai University School of Medicine, Isehara, Japan

^37^Kyoto University, Kyoto, Japan

^38^Kumamoto University, Kumamoto, Japan

^39^Miyazaki University, Miyazaki, Japan

## Methods

### Ethics statement

All experiments with hamsters were performed in accordance with the Science Council of Japan’s Guidelines for the Proper Conduct of Animal Experiments. The protocols were approved by the Institutional Animal Care and Use Committee of National University Corporation Hokkaido University (approval ID: 20-0123 and 20-0060). All protocols involving specimens from human subjects recruited at Kyoto University. All human subjects provided written informed consent. All protocols for the use of human specimens were reviewed and approved by the Institutional Review Board of Kyoto University (approval ID: R2379-3).

### Cell culture

HEK293T cells (a human embryonic kidney cell line; ATCC, CRL-3216), HEK293 cells (a human embryonic kidney cell line; ATCC, CRL-1573) and HOS-ACE2/TMPRSS2 cells (HOS cells stably expressing human ACE2 and TMPRSS2)^36,37^ were maintained in DMEM (high glucose) (Sigma-Aldrich, Cat# 6429-500ML) containing 10% fetal bovine serum (FBS, Sigma-Aldrich Cat# 172012-500ML) and 1% penicillin–streptomycin (PS) (Sigma-Aldrich, Cat# P4333-100ML). HEK293-ACE2 cells (HEK293 cells stably expressing human ACE2)^14^ were maintained in DMEM (high glucose) containing 10% FBS, 1 µg/ml puromycin (InvivoGen, Cat# ant-pr-1) and 1% PS. HEK293-ACE2/TMPRSS2 cells (HEK293 cells stably expressing human ACE2 and TMPRSS2)^14^ were maintained in DMEM (high glucose) containing 10% FBS, 1 µg/ml puromycin, 200 µg/ml hygromycin (Nacalai Tesque, Cat# 09287-84) and 1% PS. Vero cells [an African green monkey (*Chlorocebus sabaeus*) kidney cell line; JCRB Cell Bank, JCRB0111] were maintained in Eagle’s minimum essential medium (EMEM) (Sigma-Aldrich, Cat# M4655-500ML) containing 10% FBS and 1% PS. VeroE6/TMPRSS2 cells (VeroE6 cells stably expressing human TMPRSS2; JCRB Cell Bank, JCRB1819)^38^ were maintained in DMEM (low glucose) (Wako, Cat# 041-29775) containing 10% FBS, G418 (1 mg/ml; Nacalai Tesque, Cat# G8168-10ML) and 1% PS. Calu-3 cells (ATCC, HTB-55) were maintained in Eagle’s minimum essential medium (EMEM) (Sigma-Aldrich, Cat# M4655-500ML) containing 10% FBS and 1% PS. Calu-3/DSP_1-7_ cells (Calu-3 cells stably expressing DSP_1-7_)^39^ were maintained in EMEM (Wako, Cat# 056-08385) containing 20% FBS and 1% PS. Human alveolar epithelial cells derived from human induced pluripotent stem cells (iPSCs) were manufactured according to established protocols as described below (see “Preparation of human alveolar epithelial cells from human iPSCs” section) and provided by HiLung Inc. AO-ALI model was generated according to established protocols as described below (see “AO-ALI model” section). Human iPSC-derived lung organoids were generated according to established protocols as described below (see “iPSC-derived lung organoids” section). Expi293F cells (Thermo Fisher Scientific, Cat# A14527) were maintained in Expi293 expression medium (Thermo Fisher Scientific, Cat# A1435101).

### Viral genome sequencing

Viral genome sequencing was performed as previously described^18^. Briefly, the virus sequences were verified by viral RNA-sequencing analysis. Viral RNA was extracted using a QIAamp viral RNA mini kit (Qiagen, Cat# 52906). The sequencing library employed for total RNA sequencing was prepared using the NEBNext Ultra RNA Library Prep Kit for Illumina (New England Biolabs, Cat# E7530). Paired-end 76-bp sequencing was performed using a MiSeq system (Illumina) with MiSeq reagent kit v3 (Illumina, Cat# MS-102-3001). Sequencing reads were trimmed using fastp v0.21.0^40^ and subsequently mapped to the viral genome sequences of a lineage B isolate (strain Wuhan-Hu-1; GenBank accession number: NC_045512.2)^38^ using BWA-MEM v0.7.17^41^. Variant calling, filtering, and annotation were performed using SAMtools v1.9^42^ and snpEff v5.0e^43^.

### Mutation frequency calculation and phylogenetic tree reconstruction

Genomic sequences and surveillance data of 15,886,795 SARS-CoV-2 isolates were obtained from the GISAID database on August 21, 2023 (https://www.gisaid.org)^44^. The PANGO lineage of each isolate was reassigned using NextClade v2.14.0^45^. We excluded the data of SARS-CoV-2 isolate that i) was collected after August 15, 2023; ii) was isolated from non-human hosts; iii) was sampled from the original passage; and iv) whose genomic sequence is not longer than 28,000 base pairs and contains ≥2% of unknown (N) nucleotides.

We randomly selected at most 500 genomic sequences of SARS-CoV-2 in BA.1, BA.2, XBB, XBB.1, XBB.1.5, XBB.1.9, XBB.1.9.2, EG.5, EG.5.1, and EG.5.1.1 for calculating a mutation frequency (EPI SET ID: EPI_SET_231018pe). A mutation frequency of each subvariant Is calculated by dividing the number of sequences harboring the substitution of interest with the total number of sequences in that subvariant.

Next, we randomly selected at most 20 genomic sequences of SARS-CoV-2 in each XBB subvariant, resulting in 4,906 genomic sequences of SARS-CoV-2 from 248 XBB subvariants (EPI SET ID: EPI_SET_231003ue) to reconstruct a phylogenetic tree of SARS-CoV-2 in the XBB lineage. The sampled genomic sequences were aligned to the genomic sequence of Wuhan-Hu-1 SARS-CoV-2 isolate (NC_045512.2) using reference-guide multiple pairwise alignment strategy implemented in ViralMSA v1.1.24^46^. Gaps in the alignment were removed automatically using TrimAl v1.4.rev22 with-gappyout mode^47^, and the flanking edges of the alignment at positions 1–341 and 29,557–29,624 were trimmed manually. A maximum likelihood-based phylogenetic tree of representative XBB sublineages was then reconstructed from the alignment using IQ-TREE v2.2.0^48^. The best-fit nucleotide substitution model was selected automatically using ModelFinder^49^. Branch support was assessed using ultrafast bootstrap approximation^50^ with 1,000 bootstrap replicates. We omitted a genomic sequence of Wuhan-Hu-1 from the reconstructed tree and manually rooted the tree using the MRCA node including SARS-CoV-2 isolates in the original XBB subvariant.

### Reconstruction of the ancestral state of mutation

The state of having or lacking ORF9b:I5T and S:F456L substitutions was assigned to terminal nodes of the reconstructed tree based on the mutation calling data from the GISAID database. We then reconstructed the state of having ORF9b:I5T and S:F456L substitutions in the ancestral nodes from the mutation calling data obtained from the GISAID database. The reconstruction was performed using ace function of the ape R package v.5.7-1^51^ with equal-rate model. The ancestral node with a scaled likelihood of having the mutation at least 0.5 is considered having the mutation, whereas the ancestral node with the scaled likelihood less than 0.5 is considered lacking the mutation. The occurrence event of ORF9b:I5T and S:F456L substitutions was determined from the state change from lacking mutation in the ancestral node to having mutation in the adjacent descendant node. The reconstructed tree was visualized using the ggtree R package v3.8.2^52^. All the phylogenetic analyses were aided by R v4.3.1^53^.

### Modeling the relationship between amino acid substitutions and epidemic dynamics

We modeled the relationship between amino acid substitutions (not including deletions and insertions) and epidemic dynamics of SARS-CoV-2 in the XBB lineage collected in the USA from December 1, 2022 to August 15, 2023 (EPI SET ID: EPI_SET_231003vx). We used the Bayesian hierarchical multinomial logistic model described in detail in our previous study^5^. Briefly, the SARS-CoV-2 isolates were categorized into haplotypes based on their substitution profile. Substitutions observed in >200 isolates but <90% of the total isolates were selected to create the substitution profile matrix. The haplotype with <30 isolates were excluded. We also identified a cluster of co-occurring substitutions by connecting a substitution pair having Pearson’s correlation >0.9, resulting in profiles of 283 substitution clusters in 470 SARS-CoV-2 haplotypes. The representative subvariant of each haplotype was identified using the majority rule. We used an XBB.1.5 haplotype, the most abundant haplotype in the dataset, as a reference for the modeling. Finally, we counted the number of each haplotype collected in each day and created a count matrix.

Next, we applied our Bayesian hierarchical multinomial logistic model to reconstruct the relationship between amino acid substitutions and epidemic dynamics using the prepared substitution profile and count matrices. The model is 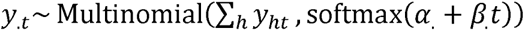 where *y_ht_* is the count of haplotype h at time *t, α_h_* and *β_h_* are intercept and slope (or growth rate) parameters for haplotype h, respectively. The slope parameter *β_h_* is derived from the Student’s *t* distribution Student’s 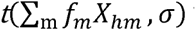 with five degrees of freedom where *f_m_* is the effect of substitution cluster *m, x_hm_* is the substitution cluster profile of haplotype *h*, and *σ* is a standard deviation. We used the Laplace distribution and half Student’s *t* distribution with five degrees of freedom as priors for *f_m_* and *σ*, respectively. The mean and standard deviation for both distributions were set to 0 and 10, respectively. Non-informative prior was set for other parameters.

The relative R_e_ of each haplotype compared to the reference haplotype or r_h_ is estimated from the equation r_h_ = exp(αβ_h_)where β_h_ is the slope parameter and y is the average viral generation time (2.1 days) (http://sonorouschocolate.com/covid19/index.php?title=Estimating_Generation_ Time_Of_Omicron). Similarly, the effect of each substitution on the relative R_e_ or *F*_h_ is calculated according to the coefficient f_h_ using the equation F_h_ = exp(αf_h_). Parameter estimation was performed by using the Markov chain Monte Carlo (MCMC) approach implemented in CmdStan v2.31.0 (https://mc-stan.org) accessed through the CmdStanr v0.5.3 R interface (https://mc-stan.org/cmdstanr/). Four independent 20,000-step MCMC chains were run including 20% of warmup iterations. We confirmed that all runs have an estimated convergence diagnostic value Rͯ is <1.01 and bulk and tail effective sampling sizes are >200, indicating the successful convergence of each run.

### Plasmid construction

Plasmids expressing the codon-optimized SARS-CoV-2 S proteins of B.1.1 (the parental D614G-bearing variant), Delta, BA.2, XBB.1, XBB.1.5, EG.5 and the two EG.5.1 derivatives were prepared in our previous studies^1,2,4,14,16,17^.

Original human ACE2 protein (residues 19–615; GenBank Accession Number NP_001358344.1) was modified to allow its efficient expression in bacteria (Escherichia coli strain BL21), but residues participating in the interaction with the SARS-CoV-2 RBD (5 Å distance) remained unaltered to keep the interaction surface identical (details about the sequence of this modified ACE2 protein are subjected to a separate publication and are available upon request; hereinafter modified ACE2 protein will be referred as mACE2). The mACE2 was inserted in pET28-14his-bdSUMO vector^54^ by restriction-free cloning and verified by sequencing.

Mammalian cell codon-optimized SARS-CoV-2 S RBDs of XBB.1 and XBB.1.5 were amplified from the expression plasmids for the codon-optimized SARS-CoV-2 S proteins of XBB.1^1^ and XBB.1.5^2^. The S RBDs of XBB.1.16 and EG.5.1 were constructed by site-directed mutagenesis with primers: XBB-K478R_R, 5’-CAG TTG GGG CCG GCC ACT CCA TTA CAT GGC CTG TTG CCA GCC TGG TAA ATC TCT G-3’ and XBB-F456L_R, 5’-GTC CCT CTC AAA TGG TTT CAG CTT GCT CTT CCT CAA CAG TCT GTA GAG GTA GTT GTA GTT GC-3’. All PCR reactions were performed by KAPA HiFi HotStart ReadyMix kit (Roche, Cat# KK2601) and subsequently assembled by yeast [*Saccharomyces cerevisiae* strain EBY100 (ATCC, Cat# MYA-4941)] homologous recombination with pJYDC1 plasmid (Addgene, Cat# 162458) as previously described^1,5,18,19,55^.

Plasmids expressing the codon-optimized SARS-CoV-2 ORF9b and ORF6 proteins were prepared in previous study (PMID: 32353859) (kindly provided by Dr. Nevan J. Krogan). SARS-CoV-2 ORF9b-based derivatives were generated by site-directed overlap extension PCR using the primers listed in **Supplementary Table S4**. The resulting PCR fragment was digested with EcoRI (New England Biolabs, Cat# R3101S) and BamHI (New England Biolabs, Cat# R3136S) and inserted into the corresponding site of the pLVX-EF1alpha-IRES-Puro vector (PMID: 32353859). Nucleotide sequences were determined by DNA sequencing services (Eurofins), and the sequence data were analyzed by Sequencher v5.1 software (Gene Codes Corporation). To generate recombinant SARS-CoV-2, the nine pmW118 plasmid vectors were subjected to amplification of the cDNA fragments (F1-F9-10) of SARS-CoV-2 EG.5.1. Nucleotide sequences were confirmed by a SeqStudio Genetic Analyzer (Thermo Fisher Scientific) and a DNA sequencing service (Fasmac).

### SARS-CoV-2 reverse genetics

Recombinant SARS-CoV-2 was generated by circular polymerase extension reaction (CPER) as previously described with modification^14,17,31,56^. In brief, 9 DNA fragments encoding the partial genome of SARS-CoV-2 were prepared by PCR using Q5 High-Fidelity DNA Polymerase (New England Biolabs, Cat# M0491S). A linker fragment encoding hepatitis delta virus ribozyme, bovine growth hormone poly A signal and cytomegalovirus promoter was also prepared by PCR with the primer-set described previously^56^. The 10 obtained DNA fragments were mixed and used for CPER.

To generate rEG.5.1-ORF9b KO and rEG.5.1-ORF9b:T5I (**Figure 7**), mutations were inserted in fragment 9 by inverse PCR-based site-directed mutagenesis with the primers listed in **Supplementary Table 4**. Nucleotide sequences were confirmed by the Sanger method as described above.

To produce recombinant SARS-CoV-2, the CPER products (25 μl) were transfected into VeroE6/TMPRSS2 cells using TransIT-X2 Dynamic Delivery System (Takara, Cat# MIR6003) according to our previous report^22^. The working virus stock was prepared from the seed virus as described below (see “SARS-CoV-2 preparation and titration” section below).

### SARS-CoV-2 preparation and titration

The working virus stocks of SARS-CoV-2 were prepared and titrated as previously described^14,17,18,57^. In this study, clinical isolates of Delta (B.1.617.2, strain TKYTK1734; GISAID ID: EPI_ISL_2378732)^16^, XBB.1.5 (strain TKYmbc30523/2022; GISAID ID: EPI_ISL_16697941)^22^, EG.5.1 (strain KU2023071028; GISAID ID: EPI_ISL_18072016), and EG.5.1.1 (strain KU2023071635; GISAID ID: EPI_ISL_18072017) were used. Also, the artificially generated recombinant viruses by the CPER technique^56^, rEG.5.1 WT, rEG.5.1 ORF9b KO, rEG.5.1 ORF9b:T5I, were used. In brief, 20 μl of the seed virus was inoculated into VeroE6/TMPRSS2 cells (5,000,000 cells in a T-75 flask). One h.p.i., the culture medium was replaced with DMEM (low glucose) (Wako, Cat# 041-29775) containing 2% FBS and 1% PS. At 3 d.p.i., the culture medium was harvested and centrifuged, and the supernatants were collected as the working virus stock.

The titer of the prepared working virus was measured as the 50% tissue culture infectious dose (TCID_50_). Briefly, one day before infection, VeroE6/TMPRSS2 cells (10,000 cells) were seeded into a 96-well plate. Serially diluted virus stocks were inoculated into the cells and incubated at 37°C for 4 d. The cells were observed under a microscope to judge the CPE appearance. The value of TCID_50_/ml was calculated with the Reed–Muench method^58^.

For verification of the sequences of SARS-CoV-2 working viruses, viral RNA was extracted from the working viruses using a QIAamp viral RNA mini kit (Qiagen, Cat# 52906) and viral genome sequences were analyzed as described above (see “Viral genome sequencing” section). Information on the unexpected substitutions detected is summarized in **Supplementary Table S5** and the raw data are deposited in the GitHub repository (https://github.com/TheSatoLab/EG.5.1).

### Yeast surface display

Yeast surface display analysis of the interaction between selected RBD variants and mACE2 (**Figure 4A**) was performed as previously described^7,8,10,16,19,34,36,40^ with some modification.

The expression of mACE2 was initiated in 800 mL of 2YT autoinducible media with trace elements (ForMedium, Cat# AIM2YT0210) until optical density achieved 0.6 cultivated at 37°C. Subsequently, expression was further induced by the addition of IPTG to 0.25 mM and continued overnight, 20°C, 220 rpm. The bacteria culture was centrifuged (5 m, 4°C, 8,000 × *g*) and the pellet was resuspended in 15 ml of a loading buffer containing 50 mM Tris-HCi (pH 8.0) and 200 mM NaCl. Cells were sonicated to extract the fused protein, centrifuged (10 m, 4°C, 3,200 × *g*) and attached on the Ni-NTA column (2 ml). The column was washed by 10 CVs of the loading buffer supplemented with 10 mM imidazole and 10 CV of PBS. 50 μg of bdSUMO protease and 1 CV of PBS were loaded into the column for the proteolysis reaction (overnight at 4°C). Finally, the column was washed with 4 CV of PBS to obtain the cleaved mACE2, given that bdSUMO protease remained attached to the Ni-column thanks to the 14 × His-tag it contained. This was proved after the elution with 4 CV of loading buffer supplemented with 300 mM imidazole and the subsequent analysis of all the loading, washing, and elution fractions by SDS-PAGE. Pure protein (**Supplementary Figure 4A**) was flash-frozen in liquid nitrogen and stored at –80°C.

Yeast expression of SARS-CoV-2 S RBD was carried out for 48 h at 20°C, and then cells were washed with PBS supplemented with bovine serum albumin at 1 g/l and incubated with 12 concentrations of mACE2 (4 pM to 10 nM, dilution series with factor 2) and 20 nM bilirubin (Sigma-Aldrich, Cat# 14370-1G). Performance comparison with Expi293F cells produced ACE2 peptidase domain (residues 18–617) was performed and is shown for XBB, XBB.1.5. and XBB.1.16. in **Supplementary Figure 4B**. RBD expression and ACE2 signal were recorded by using an automated acquisition from 96 well plates by CytoFLEX Flow Cytometer (Beckman Coulter), background binding signals were subtracted, fluorescence spill of eUnaG2 signals to red channel was compensated and data were fitted to a standard noncooperative Hill equation by nonlinear least-squares regression using Python v3.7 (https://www.python.org) as previously described^7,8,10,16,19,34,36,40^.

### SARS-CoV-2 S-based fusion assay

A SARS-CoV-2 S-based fusion assay (**Figures 4B and 4C**) was performed as previously described^1,5,14-21^. Briefly, on day 1, effector cells (i.e., S-expressing cells) and target cells (Calu-3/DSP_1-7_ cells) were prepared at a density of 0.6–0.8 × 10^6^ cells in a 6-well plate. On day 2, for the preparation of effector cells, HEK293 cells were cotransfected with the S expression plasmids (400 ng) and pDSP_8-11_ (ref.^59^) (400 ng) using TransIT-LT1 (Takara, Cat# MIR2300). On day 3 (24 h posttransfection), 16,000 effector cells were detached and reseeded into a 96-well black plate (PerkinElmer, Cat# 6005225), and target cells were reseeded at a density of 1,000,000 cells/2 ml/well in 6-well plates. On day 4 (48 h posttransfection), target cells were incubated with EnduRen live cell substrate (Promega, Cat# E6481) for 3 h and then detached, and 32,000 target cells were added to a 96-well plate with effector cells. *Renilla* luciferase activity was measured at the indicated time points using Centro XS3 LB960 (Berthhold Technologies). For measurement of the surface expression level of the S protein, effector cells were stained with rabbit anti-SARS-CoV-2 S S1/S2 polyclonal antibody (Thermo Fisher Scientific, Cat# PA5-112048, 1:100). Normal rabbit IgG (Southern Biotech, Cat# 0111-01, 1:100) was used as a negative control, and APC-conjugated goat anti-rabbit IgG polyclonal antibody (Jackson ImmunoResearch, Cat# 111-136-144, 1:50) was used as a secondary antibody. The surface expression level of S proteins (**Figure 4B**) was measured using CytoFLEX Flow Cytometer (Beckman Coulter) and the data were analyzed using FlowJo software v10.7.1 (BD Biosciences). For calculation of fusion activity, *Renilla* luciferase activity was normalized to the mean fluorescence intensity (MFI) of surface S proteins. The normalized value (i.e., *Renilla* luciferase activity per the surface S MFI) is shown as fusion activity.

### AO-ALI model

An airway organoid (AO) model was generated according to our previous report^19,60^. Briefly, normal human bronchial epithelial cells (NHBEs, Cat# CC-2540, Lonza) were used to generate AOs. NHBEs were suspended in 10 mg/ml cold Matrigel growth factor reduced basement membrane matrix (Corning, Cat# 354230). Fifty microliters of cell suspension were solidified on prewarmed cell culture-treated multiple dishes (24-well plates; Thermo Fisher Scientific, Cat# 142475) at 37°C for 10 m, and then, 500 μl of expansion medium was added to each well. AOs were cultured with AO expansion medium for 10 d. For maturation of the AOs, expanded AOs were cultured with AO differentiation medium for 5 d.

The AO-derived ALI (AO-ALI) model (**Figure 2E**) was generated according to our previous report^19,60^. For generation of AO-ALI, expanding AOs were dissociated into single cells, and then were seeded into Transwell inserts (Corning, Cat# 3413) in a 24-well plate. AO-ALI was cultured with AO differentiation medium for 5 d to promote their maturation. AO-ALI was infected with SARS-CoV-2 from the apical side.

### Preparation of human iPSC-derived alveolar epithelial cells

The ALI culture of alveolar epithelial cells (**Figure 2F**) was differentiated from human iPSC-derived lung progenitor cells as previously described^18,19,61-64^. Briefly, alveolar progenitor cells were induced stepwise from human iPSCs according to a 21-day and 4-step protocol^61^. At day 21, alveolar progenitor cells were isolated with the specific surface antigen carboxypeptidase M and seeded onto the upper chamber of a 24-well Cell Culture Insert (Falcon, #353104), followed by 7-day differentiation of alveolar epithelial cells. Alveolar differentiation medium with dexamethasone (Sigma-Aldrich, Cat# D4902), KGF (PeproTech, Cat# 100-19), 8-Br-cAMP (Biolog, Cat# B007), 3-isobutyl 1-methylxanthine (IBMX) (Fujifilm Wako, Cat# 095-03413), CHIR99021 (Axon Medchem, Cat# 1386), and SB431542 (Fujifilm Wako, Cat# 198-16543) was used for the induction of alveolar epithelial cells.

### Preparation of human iPSC-derived lung organoids

Human iPSC-derived lung organoids were used for evaluation of antiviral drugs. The iPSC line (1383D6) (provided by Dr. Masato Nakagawa, Kyoto University) was maintained on 0.5 μg/cm2 recombinant human laminin 511 E8 fragments (iMatrix-511 silk, Nippi, Cat# 892021) with StemFit AK02N medium (Ajinomoto, Cat# RCAK02N) containing 10 μM Y-27632 (FUJIFILM Wako Pure Chemical, Cat# 034-24024). For passaging, iPSC colonies were treated with TrypLE Select Enzyme (Thermo Fisher Scientific, Cat# 12563029) for 10 m at 37°C. After centrifugation, the cells were seeded onto Matrigel Growth Factor Reduced Basement Membrane (Corning, Cat# 354230)-coated cell culture plates (2.0 × 10^5^ cells/4 cm^2^) and cultured for 2 d. Lung organoids differentiation was performed in serum-free differentiation (SFD) medium of DMEM/F12 (3:1) (Thermo Fisher Scientific, Cat# 11320033) supplemented with N2 (FUJIFILM Wako Pure Chemical, Cat# 141-08941), B-27 Supplement Minus Vitamin A (Thermo Fisher Scientific, Cat# 12587001), ascorbic acid (50 μg/ml, STEMCELL Technologies, Cat# ST-72132), 1 × GlutaMAX (Thermo Fisher Scientific, Cat# 35050-079), 1% monothioglycerol (FUJIFILM Wako Pure Chemical, Cat# 195-15791), 0.05% bovine serum albumin, and 1 × PS. For definitive endoderm induction, the cells were cultured for 3 d (days 0–3) using SFD medium supplemented with 10 μM Y-27632 and 100 ng/mL recombinant Activin A (R&D Systems, Cat# 338-AC-010). For anterior foregut endoderm induction (days 3–5), the cells were cultured in SFD medium supplemented with 1.5 μM dorsomorphin dihydrochloride (FUJIFILM Wako Pure Chemical, Cat# 047-33763) and 10 μM SB431542 (FUJIFILM Wako Pure Chemical, Cat# 037-24293) for 24 h and then in SFD medium supplemented with 10 μM SB431542 and 1 μM IWP2 (REPROCELL) for another 24 h. For the induction of lung progenitors (days 5–12), the resulting anterior foregut endoderm was cultured with SFD medium supplemented with 3 μM CHIR99021 (FUJIFILM Wako Pure Chemical, Cat# 032-23104), 10 ng/ml human FGF10 (PeproTech, Cat# 100-26), 10 ng/ml human FGF7 (PeproTech, Cat# 100-19), 10 ng/ml human BMP4 (PeproTech, Cat# 120-05ET), 20 ng/ml human EGF (PeproTech, Cat# AF-100-15), and all-trans retinoic acid (ATRA, Sigma-Aldrich, Cat# R2625) for 7 d. At 12 d of differentiation, the cells were dissociated and embedded in Matrigel Growth Factor Reduced Basement Membrane to generate organoids. For lung organoid maturation (days 12–30), the cells were cultured in SFD medium containing 3 μM CHIR99021, 10 ng/ml human FGF10, 10 ng/mL human FGF7, 10 ng/ml human BMP4, and 50 nM ATRA for 8 days. At day 20 of differentiation, the lung organoids were recovered from the Matrigel, and the resulting suspension of lung organoids (small free-floating clumps) was seeded onto Matrigel-coated 96-well cell culture plates. The organoids were cultured in SFD medium containing 50 nM dexamethasone (Selleck, Cat# S1322), 0.1 mM 8-bromo-cAMP (Sigma-Aldrich, Cat# B7880), and 0.1 mM IBMX (3-isobutyl-1-methylxanthine) (FUJIFILM Wako Pure Chemical, Cat# 099-03411) for an additional 10 days before the antiviral drug experiments.

### Antiviral drug assay using SARS-CoV-2 clinical isolates and iPSC-derived lung organoids

Antiviral drug assay (**Figure 3**) was performed as previously described^29^. The human iPSC-derived lung organoids were infected with either Delta, XBB.1.5, EG.5.1, or EG.5.1.1 isolate (100 TCID_50_) at 37 °C for 2 h. The cells were washed with DMEM and cultured in DMEM supplemented with 10%LFCS, 1% PS and the serially diluted Remdesivir (Clinisciences, Cat# A17170), EIDD-1931 (an active metabolite of Molnupiravir; Cell Signalling Technology, Cat# 81178S), Ensitrelvir (MedChemExpress, Cat# HY-143216), or Nirmatrelvir (PF-07321332; MedChemExpress, Cat# HY-138687). At 72L h after infection, the culture supernatants were collected, and viral RNA was quantified using RT–qPCR (see “RT–qPCR” section below). The assay of each compound was performed in quadruplicate, and the 50% effective concentration (EC_50_) was calculated using Prism 9 software v9.1.1 (GraphPad Software).

### Airway-on-a-chip

Airway-on-a-chip (**Figures 4D and 4E**) was prepared as previously described^19,23,64^. Human lung microvascular endothelial cells (HMVEC-L) were obtained from Lonza (Cat# CC-2527) and cultured with EGM-2-MV medium (Lonza, Cat# CC-3202). For preparation of the airway-on-a-chip, first, the bottom channel of a polydimethylsiloxane (PDMS) device was precoated with fibronectin (3 μg/ml, Sigma-Aldrich, Cat# F1141). The microfluidic device was generated according to our previous report^65^. HMVEC-L cells were suspended at 5,000,000 cells/ml in EGM2-MV medium. Then, 10 μl of suspension medium was injected into the fibronectin-coated bottom channel of the PDMS device. Then, the PDMS device was turned upside down and incubated. After 1 h, the device was turned over, and the EGM2-MV medium was added into the bottom channel. After 4 d, AOs were dissociated and seeded into the top channel. Aos were generated according to our previous report^60^. AOs were dissociated into single cells and then suspended at 5,000,000 cells/ml in the AO differentiation medium. Ten microliter suspension medium was injected into the top channel. After 1 h, the AO differentiation medium was added to the top channel. In the infection experiments (**Figure 4D**), the AO differentiation medium containing either Delta, XBB.1.5, EG.5.1, or EG.5.1.1 isolate (500 TCID_50_) was inoculated into the top channel. At 2 h.p.i., the top and bottom channels were washed and cultured with AO differentiation and EGM2-MV medium, respectively. The culture supernatants were collected, and viral RNA was quantified using RT–qPCR (see “RT–qPCR” section above).

### Microfluidic device

A microfluidic device was generated according to our previous reports^23,65^. Briefly, the microfluidic device consisted of two layers of microchannels separated by a semipermeable membrane. The microchannel layers were fabricated from PDMS using a soft lithographic method. PDMS prepolymer (Dow Corning, Cat# SYLGARD 184) at a base to curing agent ratio of 10:1 was cast against a mold composed of SU-8 2150 (MicroChem, Cat# SU-8 2150) patterns formed on a silicon wafer. The cross-sectional size of the microchannels was 1 mm in width and 330 μm in height. Access holes were punched through the PDMS using a 6-mm biopsy punch (Kai Corporation, Cat# BP-L60K) to introduce solutions into the microchannels. Two PDMS layers were bonded to a PET membrane containing 3.0-μm pores (Falcon, Cat# 353091) using a thin layer of liquid PDMS prepolymer as the mortar. PDMS prepolymer was spin-coated (4,000 rpm for 60 s) onto a glass slide. Subsequently, both the top and bottom channel layers were placed on the glass slide to transfer the thin layer of PDMS prepolymer onto the embossed PDMS surfaces. The membrane was then placed onto the bottom layer and sandwiched with the top layer. The combined layers were left at room temperature for 1 d to remove air bubbles and then placed in an oven at 60°C overnight to cure the PDMS glue. The PDMS devices were sterilized by placing them under UV light for 1 h before the cell culture.

### SARS-CoV-2 infection

One day before infection, Vero cells (10,000 cells), VeroE6/TMPRSS2 cells (10,000 cells), 293-ACE2/TMPRSS2 cells (10,000 cells), and Calu-3 cells (10,000 cells) were seeded into a 96-well plate. SARS-CoV-2 [1,000 TCID_50_ for Vero cells (**Figures 2A and 7D**); 100 TCID_50_ for VeroE6/TMPRSS2 cells (**Figures 2B and 7E**); 100 TCID_50_ for 293-ACE2/TMPRSS2 cells (**Figures 2C and 7F**); and 100 TCID_50_ for Calu-3 cells (**Figures 2D and 7G**)] was inoculated and incubated at 37°C for 1 h. The infected cells were washed, and 180 µl of culture medium was added. The culture supernatant (10 µl) was harvested at the indicated timepoints and used for RT–qPCR to quantify the viral RNA copy number (see “RT–qPCR” section below). In the infection experiments using AO-ALI model (**Figures 2E and 7H**), the diluted viruses (1,000 TCID50 in 100Lμl) were inoculated onto the apical side of the culture and incubated at 37L°C for 1 h. The infected cells were washed, and 100 µl of AO differentiation medium was added. The culture supernatant (10 µl) was harvested at the indicated timepoints and used for RT–qPCR to quantify the viral RNA copy number (see “RT–qPCR” section below).

In the infection experiments using iPSC-derived alveolar epithelial cells (**Figure 2F**), working viruses were diluted with Opti-MEM (Thermo Fisher Scientific, Cat# 11058021). The diluted viruses (1,000 TCID_50_ in 100 μl) were inoculated onto the apical side of the culture and incubated at 37 °C for 1 h. The inoculated viruses were removed and washed twice with Opti-MEM. For collection of the viruses, 100 μl Opti-MEM was applied onto the apical side of the culture and incubated at 37 °C for 10 Lm. The Opti-MEM was collected and used for RT–qPCR to quantify the viral RNA copy number (see “RT–qPCR” section below). The infection experiments using an airway-on-a-chip system (**Figures 4D and 4E**) were performed as described above (see “Airway-on-a-chip” section).

### RT–qPCR

RT–qPCR was performed as previously described^14-19,29,57,64^. Briefly, 5 μl culture supernatant was mixed with 5 μl of 2 × RNA lysis buffer [2% Triton X-100 (Nacalai Tesque, Cat# 35501-15), 50 mM KCl, 100 mM Tris-HCl (pH 7.4), 40% glycerol, 0.8 U/μl recombinant RNase inhibitor (Takara, Cat# 2313B)] and incubated at room temperature for 10 m. RNase-free water (90 μl) was added, and the diluted sample (2.5 μl) was used as the template for real-time RT-PCR performed according to the manufacturer’s protocol using One Step TB Green PrimeScript PLUS RT-PCR kit (Takara, Cat# RR096A) and the following primers: Forward *N*, 5’-AGC CTC TTC TCG TTC CTC ATC AC-3’; and Reverse *N*, 5’-CCG CCA TTG CCA GCC ATT C-3’. The viral RNA copy number was standardized with a SARS-CoV-2 direct detection RT-qPCR kit (Takara, Cat# RC300A). Fluorescent signals were acquired using a QuantStudio 1 Real-Time PCR system (Thermo Fisher Scientific), QuantStudio 3 Real-Time PCR system (Thermo Fisher Scientific), QuantStudio 5 Real-Time PCR system (Thermo Fisher Scientific), StepOne Plus Real-Time PCR system (Thermo Fisher Scientific), CFX Connect Real-Time PCR Detection system (Bio-Rad), Eco Real-Time PCR System (Illumina), qTOWER3 G Real-Time System (Analytik Jena) Thermal Cycler Dice Real Time System III (Takara) or 7500 Real-Time PCR System (Thermo Fisher Scientific).

### Protein expression and purification of EG.5.1 S protein for cryo-EM

Protein expression and purification of EG.5.1 S protein were performed as previously described^66^. Briefly, the expression plasmid, pHLsec, encoding the EG.5.1LS protein ectodomain bearing six proline substitutions (F817P, A892P, A899P, A942P, K986P and V987P)^67^ and deletion of the furin cleavage site (i.e., RRAR to GSAG substitution) with a T4-foldon domain, were transfected into HEK293S GnTI(-) cells. Expressed proteins in the cell-culture supernatant were purified using a cOmplete His-Tag Purification Resin (Roche, Cat# 5893682001) affinity column, followed by Superose 6 Increase 10/300 GL size-exclusion chromatography (Cytiva, Cat# 29091596) with calcium- and magnesium-free PBS buffer.

### Cryo-EM sample preparation and data collection

The solution of EG.5.1 S protein was incubated at 37 °C for 1 h before cryo-EM grid preparation. The samples were applied to a Quantifoil R2/2 Cu 300 mesh grid (Quantifoil Micro Tools GmbH), which had been freshly glow-discharged for 60 s at 10 mA using PIB-10 (Vacuum Device). The samples were plunged into liquid ethane using a Vitrobot mark IV (Thermo Fisher Scientific) with the following settings: temperature 18°C, humidity 100%, blotting time 5 s, and blotting force 5.

Movies were collected on a Krios G4 (Thermo Fisher Scientific) operated at 300 kV with a K3 direct electron detector (Gatan) at a nominal magnification of 130,000 (0.67 Å per physical pixel), using a GIF-Biocontinuum energy filter (Gatan) with a 20 eV slit width. Each micrograph was collected with a total exposure of 1.5 s and a total dose of 50.1 e/Å^2^ over 50 frames. A total of 3,285 movies were collected at a nominal defocus range of 0.8 – 1.8 µm using EPU software (Thermo Fisher Scientific).

### Cryo-EM data processing

All datasets were processed in cryoSPARC v4.3.1^68^. Movie frames were aligned, dose-weighted, and CTF-estimated using Patch Motion correction and Patch CTF. 899,573 particles were blob-picked and reference-free 2D classification (K = 150, batch = 200, Iteration = 30) was performed to remove junk particles. 348,621 particles were used for initial model reconstruction and heterogeneous refinement. Two classes of closed states (closed-1 and closed-2) with different RBD orientations and one class of 1-up state were separated in heterogeneous refinement. The closed-1 state was processed by non-uniform refinement with C3 symmetry and CTF refinement to generate the final maps. Since the density of the RBD was unclear for the closed-2 and the 1-up states, additional processing steps were performed for these states. For the closed-2 state, once the particles were aligned with non-uniform refinement followed by aligned particles were symmetry-expanded under C3 symmetry operation. 3D classification (K = 4, force hard classification, input mode = simple) focused on the RBD without alignment was performed, and selected classes that the density of RBD was clearly resolved. A final map of closed-2 state was reconstructed with non-uniform refinement with C3 symmetry. For 1-up state, 3D classification (K = 4, force hard classification, input mode = simple) focused on the down RBD and up RBD without alignment was performed, and selected classes that the density of up RBD was clearly resolved. A final map of 1-up state was reconstructed with non-uniform refinement with C1 symmetry. C1 for 1-up state after removing duplicate particles. To support model building, a local refinement focusing on down RBD in closed-2 and down and up RBD in 1-up states was carried out.

The reported global resolutions are based on the gold-standard Fourier shell correlation curves (FSC = 0.143) criteria. Local resolutions were calculated with cryoSPARC^69^. Workflows of data processing were shown in **Supplementary Figure 1A**. Figures related to data processing and reconstructed maps were prepared with UCSF Chimera v1.17.1^70^ and UCSF Chimera X v1.6.1^71^.

### Cryo-EM model building and analysis

Structures of SARS-CoV-2 XBB.1 S protein closed-1 state (PDB: 8IOS^1^) or closed-2 state (PDB: 8IOT) were fitted to the corresponding maps using UCSF Chimera. Iterative rounds of manual fitting in Coot v0.9.6^72^ and real-space refinement in Phenix v1.20^73^ were carried out to improve non-ideal rotamers, bond angles, and Ramachandran outliers. The final model was validated with MolProbity^74^. The structure models shown in surface, ribbon and stick presentation in figures were prepared with PyMOL v2.5.0 (http://pymol.sourceforge.net).

### Animal experiments

Animal experiments (**Figure 6** and **Supplementary Figure 2**) were performed as previously described^1,5,15-19,22^. Syrian hamsters (male, 4 weeks old) were purchased from Japan SLC Inc. (Shizuoka, Japan). For the virus infection experiments, hamsters were anesthetized by intramuscular injection of a mixture of 0.15 mg/kg medetomidine hydrochloride (Domitor^®^, Nippon Zenyaku Kogyo), 2.0 mg/kg midazolam (Dormicum^®^, FUJIFILM WAKO, Cat# 135-13791) and 2.5 mg/kg butorphanol (Vetorphale^®^, Meiji Seika Pharma) or 0.15 mg/kg medetomidine hydrochloride, 4.0 mg/kg alphaxaone (Alfaxan^®^, Jurox) and 2.5 mg/kg butorphanol. EG.5.1, EG.5.1.1, XBB1.5, Delta (10,000 TCID_50_ in 100 µl), or saline (100 µl) were intranasally inoculated under anesthesia. Oral swabs were collected at the indicated timepoints. Body weight was recorded daily by 7 d.p.i. Enhanced pause (Penh), the ratio of time to peak expiratory follow relative to the total expiratory time (Rpef) were measured every day until 7 d.p.i. of the EG.5.1-, EG.5.1.1-, XBB.1.5-, and Delta-infected hamsters (see below). Lung tissues were anatomically collected at 2 and 5 d.p.i. The viral RNA load in the oral swabs and respiratory tissues was determined by RT–qPCR. These tissues were also used for IHC and histopathological analyses (see below).

### Lung function test

Lung function tests (**Figure 6A**) were routinely performed as previously described^1,5,15,17-19,22^. The two respiratory parameters (Penh and Rpef) were measured by using a Buxco Small Animal Whole Body Plethysmography system (DSI) according to the manufacturer’s instructions. In brief, a hamster was placed in an unrestrained plethysmography chamber and allowed to acclimatize for 30 s. Then, data were acquired over a 2.5-minute period by using FinePointe Station and Review software v2.9.2.12849 (DSI).

### Immunohistochemistry

Immunohistochemistry (IHC) (**Figure 6C** and **Supplementary Figure 2**) was performed as previously described^1,5,15-19,22^ using an Autostainer Link 48 (Dako). The deparaffinized sections were exposed to EnVision FLEX target retrieval solution high pH (Agilent, Cat# K8004) for 20 m at 97°C for activation, and a mouse anti-SARS-CoV-2 N monoclonal antibody (clone 1035111, R&D Systems, Cat# MAB10474-SP, 1:400) was used as a primary antibody. The sections were sensitized using EnVision FLEX for 15 m and visualized by peroxidase-based enzymatic reaction with 3,3′-diaminobenzidine tetrahydrochloride (Dako, Cat# DM827) as substrate for 5 m. The N protein positivity was evaluated by certificated pathologists as previously described. Images were incorporated as virtual slides by NDP.scan software v3.2.4 (Hamamatsu Photonics). The N-protein positivity was measured as the area using Fiji software v2.2.0 (ImageJ).

### H&E staining

H&E staining (**Figure 6D** and **Supplementary Figure 3**) was performed as previously described^1,5,15-19,22^. Briefly, excised animal tissues were fixed with 10% formalin neutral buffer solution and processed for paraffin embedding. The paraffin blocks were sectioned at a thickness of 3 µm and then mounted on MAS-GP-coated glass slides (Matsunami Glass, Cat# S9901). H&E staining was performed according to a standard protocol.

### Histopathological scoring

Histopathological scoring (**Figure 6E**) was performed as previously described^1,5,15-19,22^. Pathological features, including (i) bronchitis or bronchiolitis, (ii) hemorrhage with congestive edema, (iii) alveolar damage with epithelial apoptosis and macrophage infiltration, (iv) hyperplasia of type II pneumocytes, and (v) the area of hyperplasia of large type II pneumocytes, were evaluated in each hamsters by certified pathologists, and the degree of these pathological findings was arbitrarily scored using a four-tiered system as 0 (negative), 1 (weak), 2 (moderate), and 3 (severe). The “large type II pneumocytes” are type II pneumocytes with hyperplasia exhibiting more than 10-μm-diameter nuclei. We described “large type II pneumocytes” as one of the notable histopathological features of SARS-CoV-2 infection in our previous studies. The total histological score is the sum of these five indices.

### Transfection, western blotting, SeV Infection and reporter assay

HEK293 cells were transfected using PEI Max (Polysciences) according to the manufacturer’s protocol. For Western blotting, cells (in 12 well) were cotransfected with the pLVX-EF1alpha-IRES-Puro-based 2×Strep-tagged expression plasmids (12.5, 50, 200 or 800 ng for **Figure 7B**; 300, 600 or 900 ng for **Figure 7D**) together with an empty vector (normalized to 1 μg per well). For luciferase reporter assay, cells (in 96 well) were cotransfected with 10 ng of either p125Luc (expressing firefly luciferase driven by human IFNB1 promoter; kindly provided by Dr. Takashi Fujita)^75^ and the pLVX-EF1alpha-IRES-Puro-based 2×Strep-tagged expression plasmids (1.25, 5, 20 or 80 ng for Figures 7B; 30, 60 or 90 ng for **Figure 7D**). The amounts of transfected plasmids were normalized to 100 ng per well. At 24 h post transfection, SeV (strain Cantell, clone cCdi; GenBank accession no. AB855654)^76^ was inoculated into the transfected cells at multiplicity of infection (MOI) 100.

The luciferase reporter assay was performed 24 h.p.i. as previously described^28,77^. Briefly, 50 μl cell lysate was applied to a 96-well plate (Nunc), and the firefly luciferase activity was measured using a PicaGene BrillianStar-LT luciferase assay system (Toyo-b-net), and the input for the luciferase assay was normalized by using a CellTiter-Glo 2.0 assay kit (Promega) following the manufacturers’ instructions. For this assay, a GloMax Explorer Multimode Microplate Reader 3500 (Promega) was used.

Western Blotting was performed as previously described^28,77^. Briefly, transfected cells were lysed with RIPA buffer (25 mM HEPES [pH 7.4], 50 mM NaCl, 1 mM MgCl2, 50 mM ZnCl2, 10% glycerol, 1% Triton X-100) containing a protease inhibitor cocktail (Roche). For blotting, anti-Strep-tag II antibody (Abcam, Cat# ab76949) and anti-α-Tubulin antibody (Sigma, Cat# T9026) were used as primary antibody. Horseradish peroxidase-conjugated anti-mouse IgG antibody (Cell Signaling, Cat# 7076) and Horseradish peroxidase-conjugated anti-rabbit IgG antibody (Cell Signaling, Cat# 7074) were used as secondary antibody.

### Statistics and reproducibility

Statistical significance was tested using a two-sided Mann–Whitney *U* test, a two-sided Student’s *t* test, a two-sided Welch’s *t* test, or a two-sided paired *t-*test unless otherwise noted. The tests above were performed using Prism 9 software v9.1.1 (GraphPad Software).

In the time-course experiments (**Figure 2A–F, 4C–D, 6A–B, E, 6D–H, 7D-H**), a multiple regression analysis including experimental conditions (i.e., the types of infected viruses) as explanatory variables and timepoints as qualitative control variables was performed to evaluate the difference between experimental conditions thorough all timepoints. The initial time point was removed from the analysis. The *P* value was calculated by a two-sided Wald test. Subsequently, familywise error rates (FWERs) were calculated by the Holm method. These analyses were performed on R v4.2.1 (https://www.r-project.org/).

In **Figure 4C–D, and Supplementary Figure 1**, photographs shown are the representative areas of at least two independent experiments by using four hamsters at each timepoint.

## Data availability

Surveillance datasets of SARS-CoV-2 isolates are available from the GISAID database (https://www.gisaid.org; EPI_SET_231018pe, EPI_SET_231003ue, and EPI_SET_231003vx). The supplemental table for each GISAID dataset is available in the GitHub repository (https://github.com/TheSatoLab/EG.5.1). The atomic coordinates and cryo-EM maps for the structures of the EG.5.1LS protein alone closed state 1 (8WMF, EMD-37651), closed state 2 (8WMD, EMD-37650), 1-up (EMD-37648) are available in the Protein Data Bank (www.rcsb.org) and Electron Microscopy Data Bank (www.ebi.ac.uk/emdb/).

## Code availability

The computational codes used in the present study are available in the GitHub repository (https://github.com/TheSatoLab/EG.5.1).

## Supplementary files

Supplementary Table 1. Effect of amino acid substitution in the 12 SARS-CoV-2 proteins on R_e_ and relating modeling parameters of SARS-CoV-2 in the XBB lineage circulated in the USA from December 1, 2022 to September 15, 2023.

Supplementary Table 2. Estimated relative R_e_ and modeling parameters of haplotypes of SARS-CoV-2 in the XBB lineage circulated in the USA from December 1, 2022 to September 15, 2023.

Supplementary Table 3. Cryo-EM data collection, refinement and validation statistics

Supplementary Table 4. Primers used in this study for preparation of SARS-CoV-2 ORF9b expression plasmid

Supplementary Table 5. Summary of unexpected amino acid mutations detected in the working virus stocks

**Supplementary Figure 1. Workflow of cryo-EM data processing for EG.5.1 S and structural comparison for EG.5.1 and XBB.1.5 S, related to Figure 5**

(A) (Left) Representative micrograph (scale bars, 50 nm) and 2D class images. (Right) Cryo-EM data processing flowchart for EG.5.1 S. (B) Global resolution assessment of cryo-EM maps by gold-standard Fourier shell correlation (FSC) curves at the 0.143 criteria. The calculated values of local resolution was colored at grid point of cryo-EM maps. (C) Superimposed RBD structures of EG.5.1 S closed-1 and closed-2. An arrow indicates 370-375 residues of RBD that show different loop structure in these two closed states. (D) Superimposed amino-acid residues that are substituted in closed-1 state of spike protein in EG.5.1 (red) as compared to XBB.1.5 (cyan). (E)I The models fit to corresponding cryo-EM maps at Q52H and F456L substitution. Arrows indicate the substituted amino-acid residues, Q52H and F456L, between EG.5.1 and XBB.1.5.

**Supplementary Figure 2. Distribution of SARS-CoV-2 N-positive cells in the lungs of infected hamsters, related to Figure 6**

N-positive area in the lungs of infected hamsters at 2 d.p.i (**A**) and 5 d.p.i (**B**) (4 hamsters per infection group). N-protein immunohistochemistry(top) and the digitalized N-positive area (bottom, indicated in red) are shown. The red numbers in the bottom panels indicate the percentage of N-positive area. Summarized data are shown in a bar graph (right). Representative images are shown in Figure 6C.

**Supplementary Figure 3. Histological observations in infected hamsters, related to Figure 6**

Type II pneumocytes in the lungs of infected hamsters at 2 d.p.i. (**A**) and 5 d.p.i (**B**) (4 hamsters per infection group). H&E staining (top) and the digitalized inflammatory area with type II pneumocytes (bottom, indicated in red) are shown. The red numbers in the bottom panels indicate the percentage of inflammatory area with type II pneumocytes. Summarized data are shown in a bar graph (**right**). Representative images are shown in Figure 6D.

**Figure S4. Protein-engineered mACE2 protein, related to Figure 4**

(**A**) mACE2 protein isolated after one purification step on-column cleavage by bdSUMO-protease. Molecular size marker is Flash Protein Ladder, FPL-008, Gel Company, USA.

(**B**) Comparison between mACE2 and Expi293F cells produced ACE2 peptidase domain shows tighter interactions with the mACE2 despite the intact binding site. Notably, the effect of mutations in different RBDs is similar between ACE2-WT and mACE2.

