## Supplementary material for "Virological characteristics of the SARS-CoV-2 Omicron EG.5.1 variant": Table S3

**Supplementary Table 3. Cryo-EM data collection, refinement and validation statistics**

|  | **SARS-CoV-2 EG.5.1 spike** | | | |
| --- | --- | --- | --- | --- |
| **Data collection and processing** | **closed-1** | **closed-2** | | **1-up** |
| EMDB ID | EMD-37651 | EMD-37650 | | EMD-37648 |
| PDB ID | 8WMF | 8WMD | | - |
| Microscope | Krios G4 | | | |
| Camera  energy filter | Gatan K3  Gatan Biocontinuum | | | |
| slit width | 20 | | | |
| Magnification | 130,000 | | | |
| Recording mode | counting | | | |
| Voltage (kV) | 300 | | | |
| Electron exposure (e–/Å^2^) | 50.1 | | | |
| Exposure time (s) | 1.5 | | | |
| Number of raw frames | 50 | | | |
| Defocus range (μm) | -0.8 to -1.8 | | | |
| Pixel size (Å) | 0.67 | | | |
| Initial particle  images (no.) | 899,573 | | | |
| Final particle  images (no.) | 88,644 | 43,647 | | 22,926 |
| Symmetry imposed | C3 | C3 | | C1 |
| Map resolution (Å)  FSC 0.143 | 2.51 | 2.89 | | 3.34 |
| **Refinement** |  | |  | |
| Initial model used  (PDB code) | 8IOS | 8IOT | | - |
| Model composition  Protein residues  Ligands | 1074  NAG:19 | 1021  NAG:11 | | -  - |
| Map CC | 0.86 | 0.82 | | - |
| R.m.s. deviations  Bond lengths (Å)  Bond angles (°) | 0.002  0.455 | 0.004  0.915 | | -  - |
| Validation  MolProbity score  Clashscore  Rotamer outliers (%) | 1.44  3.45  0.00 | 1.51  4.78  0.00 | | -  -  - |
| Ramachandran plot  Favored (%)  Allowed (%)  Outliers (%) | 95.66  4.34  0.00 | 96.12  3.88  0.00 | | -  -  - |
