## Supplementary figures and images for "Virological characteristics of the SARS-CoV-2 Omicron EG.5.1 variant"

### Figure S1

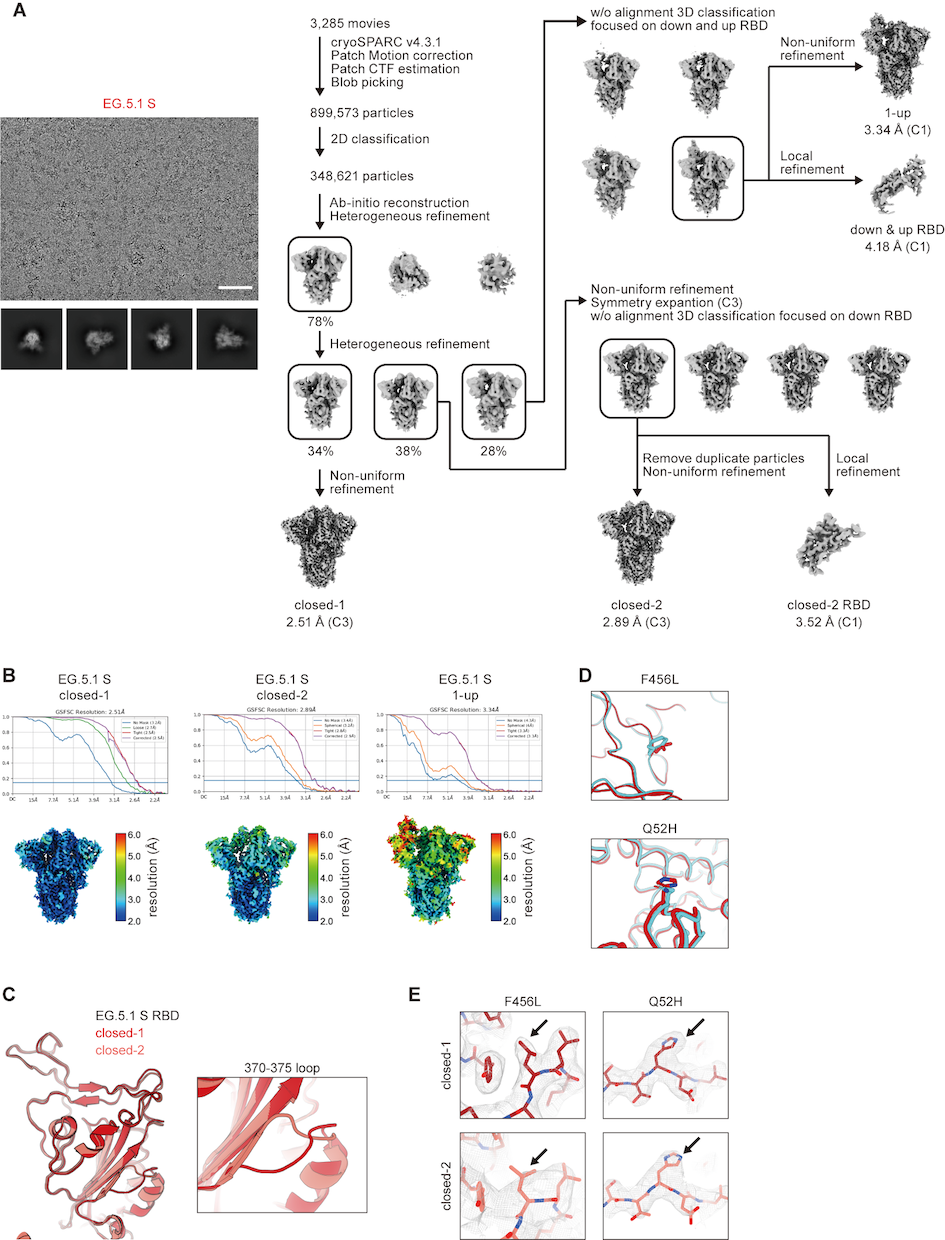

### Figure S2

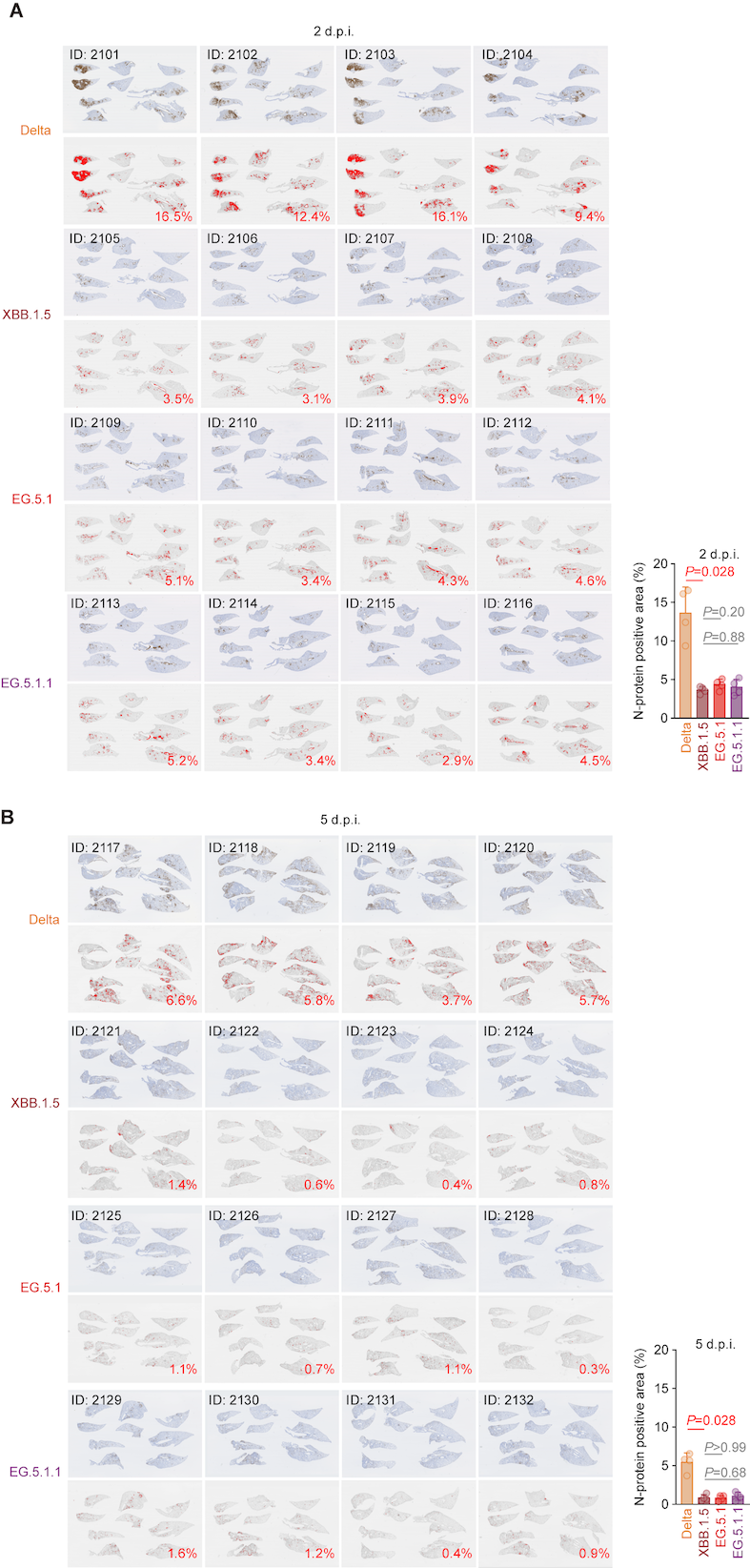

### Figure S3

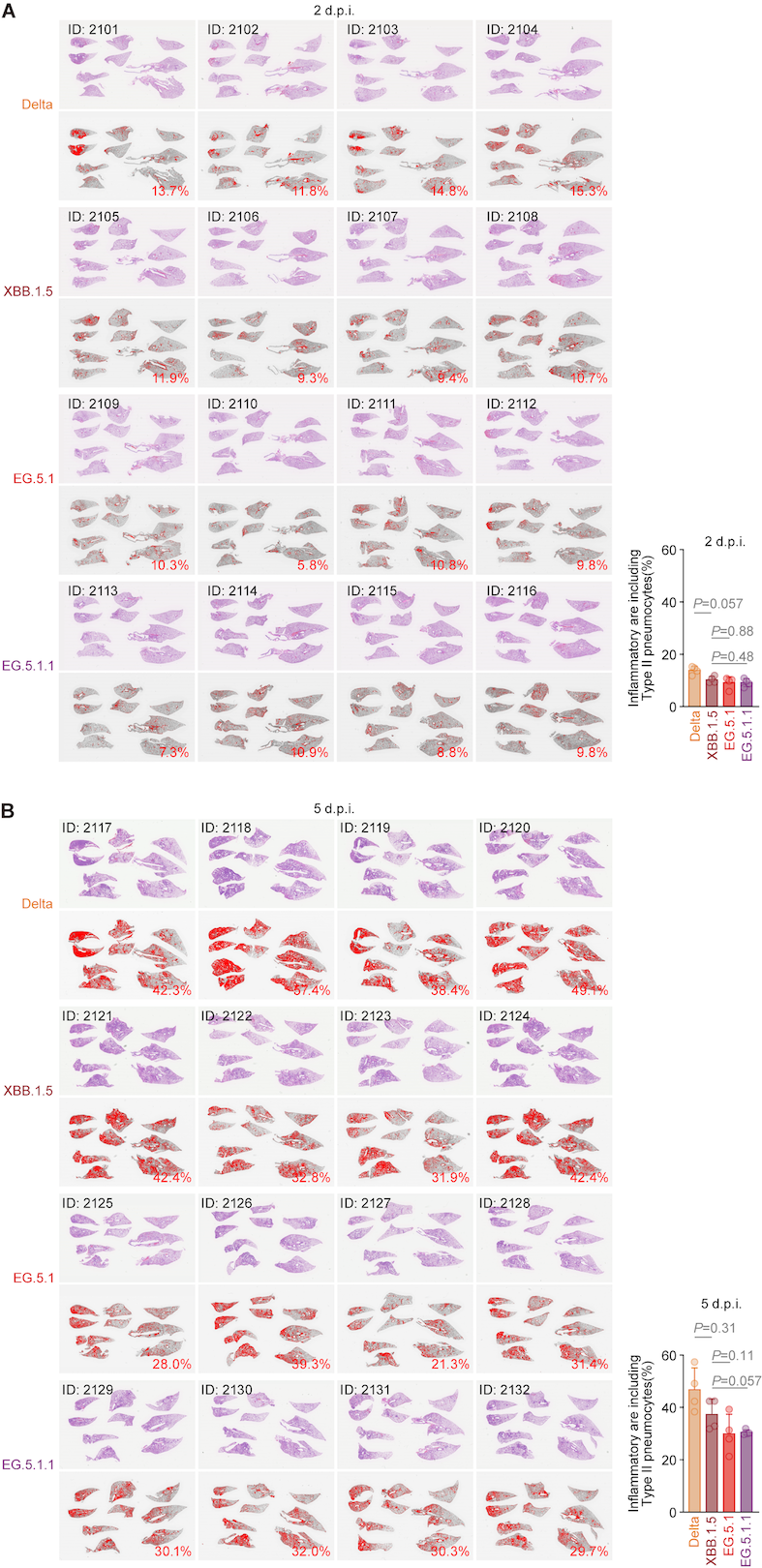

### Figure S4

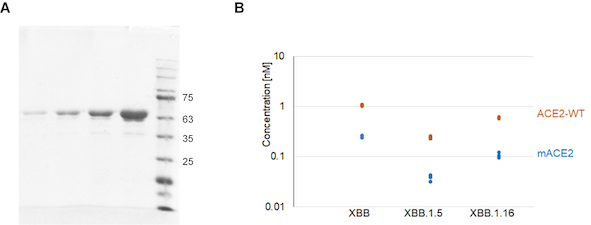
